# LGR5-high colorectal cancer defines a Wnt-active stem-like state with multimodal therapeutic vulnerabilities

**DOI:** 10.64898/2026.08.26.747295

**Authors:** Lucía Paniagua-Herranz, Alejandro Feito, Cristian Privat, Luis Álvarez-Carrión, Bernard Doger, Andrés R. Tejedor, Juan Antonio Ardura, Verónica Alonso, Cristina Nieto-Jiménez, Carlos Alonso-Moreno, Víctor Moreno, Emiliano Calvo, Balazs Gyorffy, Jorge R. Espinosa, Alberto Ocana

**Affiliations:** Experimental Therapeutics CRIS Cancer Unit, Hospital Clinico San Carlos and IdSCC and CIBERONC, Madrid, Spain; Department of Physical Chemistry, Universidad Complutense de Madrid, Av. Complutense s/n, Madrid 28040, Spain; Instituto Pluridisciplinar, Universidad Complutense de Madrid, P.° de Juan XXIII, 1, Moncloa - Aravaca, 28040 Madrid, Spain; CANMET research group, Cátedra INTHEOS-START-CEU de Oncología de precision, Departamento de Ciencias Médicas Básicas, Facultad de Medicina, Instituto de Medicina Molecular Aplicada (IMMA), Universidad San Pablo-CEU, CEU Universities ES, Urbanización Montepríncipe, 28660 Boadilla del Monte, Spain; START Madrid-FJD, Hospital Fundación Jiménez Díaz, Madrid, Spain; Yusuf Hamied Department of Chemistry, University of Cambridge, Lensfield Road, Cambridge CB2 1EW, United Kingdom; Bone Physiopathology Laboratory, Cátedra INTHEOS-START-CEU de Oncología de precisión, Departamento de Ciencias Médicas Básicas, Facultad de Medicina, Instituto de Medicina Molecular Aplicada (IMMA), Universidad San Pablo-CEU, CEU Universities, Urbanización Montepríncipe, 28660 Boadilla del Monte, Madrid, Spain; Universidad de Castilla-La Mancha, Departamento de Química Inorgánica, orgánica y bioquímica. Facultad de Farmacia-Centro de Innovación en Química Avanzada (ORFEO-CINQA), Unidad nanoDrug, 02008 Albacete, Spain; START Madrid-CIOCC, Centro Integral Oncológico Clara Campal, 28050 Madrid, Spain; Department of Bioinformatics, Semmelweis University, H-1094, Budapest, Hungary; Institute of Transdisciplinary Discoveries, Medical School, University of Pecs, H-7624, Pecs, Hungary; Cancer Biomarker Research Group, Institute of Molecular Life Sciences, HUN-REN Research Centre for Natural Sciences, H-1117, Budapest, Hungary; PhAsIca BioScience SL. Calle Velázquez 27, 1° exterior derecha, Madrid, 28001, Spain; Cátedra INTHEOS-START-CEU. START Madrid-FJD, Hospital Fundación Jiménez Díaz, Madrid, Spain

**Keywords:** Colorectal cancer · LGR5 · Druggability · Biomolecular condensates · Antibody-drug conjugates

## Abstract

Leucine-rich repeat-containing G protein-coupled receptor 5 (LGR5) marks a stem-like colorectal cancer population but is difficult to target directly because of its expression in normal intestinal stem cells. Here, we integrated transcriptomic, somatic mutation, and copy-number data from 396 colorectal tumors to define the LGR5-high molecular state and identify therapeutic vulnerabilities through modality-matched tractability analysis. LGR5-high tumors displayed a coordinated Wnt-active, stem-like transcriptional program associated with extracellular matrix remodeling, APC mutations, and IGF2 amplification, yet lacked selective genetic dependencies among the identified genes. We therefore assessed candidate targets across three complementary dimensions: surface accessibility, structural ligandability, and biomolecular condensate behavior. ENPP3 emerged as the strongest candidate for antibody-based delivery, combining tumor-associated expression, an accessible ectodomain, internalization capacity, and clinical antibody-drug conjugate precedent. PLCB4 showed the strongest structural ligandability, with independent algorithms identifying its catalytic site and additional predicted cavities. NKD1 lacked conventional ligandable pockets but formed RNA-stabilized condensates and partitioned into DVL2–AXIN1 assemblies, reducing AXIN1 self-association and DVL2–AXIN1 contacts in molecular simulations. Together, these findings define LGR5-high colorectal cancer as a therapeutically tractable molecular state and establish a modality-matched framework for translating tumor-associated signatures into experimentally testable therapeutic hypotheses.

## Background

Colorectal cancer (CRC) remains a major cause of cancer mortality worldwide (Sung et al. 2021), driven largely by the development of metastatic disease despite advances in screening and systemic therapy (Siegel et al. 2022). Its marked molecular heterogeneity continues to limit the identification of therapeutically actionable tumor states, underscoring the need to define biologically coherent subsets that can be matched to targeted interventions.

LGR5 (leucine-rich repeat-containing G protein-coupled receptor 5) is a canonical Wnt/β-catenin target and marker of intestinal stem cells (Barker et al. 2007). In CRC, LGR5-positive cells define tumor-initiating populations with enhanced self-renewal, therapeutic resistance, and metastatic capacity (Schepers et al. 2012; Merlos-Suárez et al. 2011), and elevated LGR5 expression is associated with adverse clinical outcome (Jiang et al. 2013). LGR5 reinforces Wnt signaling through R-spondin-dependent inhibition of the E3 ubiquitin ligases RNF43 and ZNRF3, thereby sustaining β-catenin-driven stemness programs (de Lau et al. 2011). Direct targeting, however, is constrained by LGR5 expression in normal intestinal stem cells, limiting the therapeutic window of monospecific approaches.

Genomic profiling has defined the transcriptomic, mutational, and copy-number architecture of CRC (Muzny et al. 2012), yet the molecular state associated with high LGR5 expression and its therapeutic implications remain incompletely resolved (Guinney et al. 2015). Addressing this gap requires moving beyond molecular description to determine which components of this state represent actionable vulnerabilities and which therapeutic modality is best suited to engage them.

Antibody-based therapies, including antibody–drug conjugates, require surface accessibility, favorable epitope topology, and efficient internalization and payload delivery (Tsuchikama and An 2018; Ritchie, Tchistiakova, and Scott 2013; López de Sá et al. 2023), whereas small-molecule strategies depend on structurally ligandable cavities that can increasingly be identified from predicted protein structures (Bertoni et al. 2026; Aggarwal et al. 2022; Le Guilloux, Schmidtke, and Tuffery 2009). Pocket detection alone, however, does not establish druggability (Schmidtke and Barril 2010; Scardino, Filippo, and Cavasotto 2023), and many proteins remain inaccessible to either modality. Biomolecular condensates provide a complementary axis of therapeutic tractability (Mitrea et al. 2022), particularly relevant to Wnt signaling, which is organized through phase-separated assemblies at opposing pathway states: the Axin-scaffolded destruction complex in the Wnt-off state (Nong et al. 2021) and DVL2-driven receptor signalosomes following pathway activation (Kang et al. 2022).

Here, we define the multi-omic landscape of LGR5-high CRC and systematically map its molecular components across three complementary dimensions of therapeutic tractability. LGR5-high tumors exhibit a coordinated Wnt-active, stem-like program associated with extracellular matrix remodeling and a distinct genomic context, yet lack selective genetic dependencies. Modality-matched prioritization identifies ENPP3 as the strongest candidate for antibody-based delivery, PLCB4 as the most ligandable intracellular target, and NKD1 as a conventionally inaccessible protein with intrinsic condensate-forming capacity. Coarse-grained simulations indicate that NKD1 co-condenses with AXIN1–DVL2 assemblies and remodels their interaction network, providing a candidate biophysical basis for NKD1’s reported role in Wnt feedback regulation. Together, these findings outline a framework for converting tumor-associated molecular states into modality-matched therapeutic hypotheses, extending it to candidates that fall outside the reach of conventional antibody-or pocket-directed approaches

## Methods

### Data source and study cohort

Genomic and transcriptomic data were obtained from The Cancer Genome Atlas (TCGA) colorectal adenocarcinoma cohorts (COAD/READ; n = 396). RNA-seq read counts, somatic mutation calls, and copy-number variation (CNV) data were retrieved from matched primary tumor samples and integrated for downstream analyses.

### LGR5-based tumor stratification

Tumors were stratified according to LGR5 expression using RNA-seq read counts. Samples with LGR5 expression ≥500 counts were classified as LGR5-high, and those below this threshold as LGR5-low. The cutoff was empirically defined to distinguish tumors with high LGR5 expression from those with low or background expression.

### Differential gene expression

Transcriptome-wide expression was compared between LGR5-high and LGR5-low tumors using the Mann–Whitney U test. Genes were considered enriched in LGR5-high tumors when they met all predefined criteria: p ≤ 0.001, fold change (FC) ≥1.5, and expression ≥500 RNA-seq counts in the LGR5-high group.

### Somatic mutation analysis

To identify somatic alterations associated with LGR5 expression, tumors carrying a given mutation were compared with wild-type tumors using the Mann–Whitney U test. Analyses were restricted to genes mutated in ≥5 tumors. Mutations were considered associated with increased LGR5 expression when mutant tumors showed FC ≥1.44 relative to wild-type tumors and p ≤ 0.05.

### Copy-Number Variation Analysis

Copy-number alterations were analyzed to identify genomic events associated with LGR5 expression. Amplifications and deletions were evaluated independently by comparing LGR5 expression in tumors harboring each alteration with that in copy-number-neutral tumors using the Mann–Whitney U test. Analyses were restricted to events present in ≥15 tumors. Amplifications were considered associated with increased LGR5 expression when they met predefined thresholds of p ≤ 0.05 and fold change (FC) ≥1.44, whereas deletions meeting p ≤ 0.05 were retained for analysis. Amplification and deletion results were subsequently integrated to define the copy-number landscape associated with LGR5 expression.

### Gene Expression Analysis

RNA-seq data were analyzed to assess the expression of LGR5 and candidate surface and non-surface protein-coding genes across tumor and normal tissues. Expression data from The Cancer Genome Atlas (TCGA) and Genotype-Tissue Expression (GTEx) projects were accessed through Gene Expression Profiling Interactive Analysis 2 (GEPIA2; http://gepia2.cancer-pku.cn/; last accessed July 17, 2026) (Tang et al. 2019; Weinstein et al. 2013; Lonsdale et al. 2013). Tumor and corresponding normal-tissue expression profiles were compared to assess tissue distribution and tumor-associated enrichment of candidate targets.

### CRISPR Dependency Analysis

CRISPR-Cas9 dependency data for genes identified as overexpressed, mutated, or copy-number altered in LGR5-high COAD tumors were obtained from the Cancer Dependency Map (DepMap; https://depmap.org/portal/; last accessed July 17, 2026) (DepMap, Broad 2025; Arafeh et al. 2025; Meyers et al. 2017; Shi et al. 2021). Analyses were restricted to COAD cell lines with high LGR5 expression (log2(TPM+1) ≥ 5) . Gene dependency was assessed using CERES scores, which estimate the effect of gene knockout on cancer-cell fitness while accounting for copy-number-related biases in CRISPR screening. More negative CERES scores indicate greater dependency, with scores <−0.5 considered indicative of a substantial impairment in cell proliferation or survival following gene knockout.

### Surfaceome Analysis

The subcellular localization of genes overexpressed in LGR5-high COAD tumors was assessed using the in silico human Surfaceome database (https://wlab.ethz.ch/surfaceome/; last accessed July 17, 2026) (Bausch-Fluck et al. 2018). Proteins annotated as cell-surface localized were subsequently evaluated as potential candidates for antibody-based therapeutic targeting.

### Structure-Based Pocket Detection and Ligandability Analysis

Three-dimensional protein structures were analyzed to identify potential ligand-binding cavities and assess structural ligandability. Because experimentally determined structures were unavailable for most candidates, predicted structures were obtained from the AlphaFold Protein Structure Database (https://alphafold.ebi.ac.uk/) (Bertoni et al. 2026). AlphaFold models were analyzed for SMOC2 (AF-Q9H3U7-F1), ENPP3 (AF-O14638-F1), CAPN6 (AF-Q9Y6Q1-F1), APOLD1 (AF-Q96LR9-F1), NKD1 (AF-Q969G9-F1), SATB1 (AF-Q01826-F1), MEX3A (AF-A1L020-F1), PGAP1 (AF-Q75T13-F1), GDPD5 (AF-Q8WTR4-F1), COL9A3 (AF-Q14050-F1), SLC6A6 (AF-P31641-F1), PLEKHB1 (AF-Q9UF11-F1), PLCB4 (AF-Q15147-F1), ITPR2 (AF-Q14571-2-F1), and FREM1 (AF-Q5H8C1-F1). H19 was excluded because it encodes a long non-coding RNA. The available AlphaFold model for ITPR2 provided incomplete sequence coverage, which was considered when interpreting pocket predictions.

Potential ligand-binding sites were independently evaluated using three complementary approaches: P2Rank implemented through the PrankWeb server (https://prankweb.cz/) (Polák et al. 2025; Krivák and Hoksza 2018), DeepPocket (Aggarwal et al. 2022), and DoGSiteScorer (Volkamer et al. 2010, 2011, 2012) implemented through the ProteinsPlus platform (https://proteins.plus/) (Ehrt et al. 2025). For PrankWeb, default parameters were used, including evolutionary conservation analysis with HMMER (Eddy 2011). DeepPocket analyses used the classification model first_model_fold1_best_test_auc_85001.pth.tar and segmentation model seg0_best_test_IOU_91.pth.tar, as described in the original publication. The number of pockets subjected to segmentation was set to 10, with all remaining parameters maintained at default values. DoGSiteScorer was run using default parameters, including subpocket detection and druggability assessment.

Because these algorithms use distinct scoring systems and do not provide directly equivalent thresholds for pocket confidence or druggability, predefined study-specific criteria were used to prioritize candidate cavities (Paiva et al. 2026; Zillmer and Walther 2025). For P2Rank, pockets with scores ≥10.0 and probabilities ≥0.5 were retained; scores of 10.0–15.0 with probabilities of 0.5–0.6 were classified as moderate-confidence, with values exceeding these thresholds considered higher-confidence. For DeepPocket, cavities with CNNconfidence ≥0.5 and druggability ≥0.5 were retained, with values of 0.5–0.7 and 0.5–0.6, respectively, classified as moderate-confidence. For DoGSiteScorer, pockets with SimpleScore ≥0.6 and DrugScore ≥0.6 were retained, with values of 0.6–0.7 for both metrics classified as moderate-confidence. Cavities below these predefined thresholds were not prioritized. Concordance across independent algorithms, together with structural context and functional annotation, was used to prioritize targets for subsequent tractability assessment.

### Prediction of Saturation Concentration (Csat)

Intrinsic propensity for homotypic phase separation was assessed using a machine-learning (ML) model trained on coarse-grained molecular dynamics simulations performed with the CALVADOS2 force field (von Bülow et al. 2025). The model predicts transfer free energies between dilute and condensed phases and estimates the corresponding saturation concentration (Csat) at 293 K and 150 mM NaCl, with lower Csat values indicating greater intrinsic propensity for phase separation.

Intrinsically disordered regions (IDRs) were defined from AlphaFold Protein Structure Database models using a pLDDT threshold <70 (Alvarez et al. 2026). Proteins with ≥40% of their sequence classified as disordered were selected for Csat analysis. For each eligible protein, predictions were performed using both the concatenated IDR sequence and the full-length protein sequence to assess the contribution of sequence context to predicted phase-separation propensity.

### Molecular Dynamics Simulations

Coarse-grained molecular dynamics simulations were performed using the residue-resolution Mpipi-Recharged force field (R. Tejedor et al. 2025), in which each amino acid is represented by a single interaction site. Hydrophobic interactions were modeled using a Wang–Frenkel potential with a cutoff of 3σ, whereas electrostatic interactions between charged residues were described by a Yukawa potential with a 3.5-nm cutoff. The Debye screening length was calculated self-consistently for 150 mM NaCl using the temperature-dependent dielectric constant. Interactions involving structured regions were scaled to account for buried residues, using factors of 0.7 for structured–structured interactions and √0.7 for structured–disordered interactions. Bonded interactions were described by a harmonic potential (k = 9.6 kcal·mol⁻¹·Å⁻²; r₀ = 3.81 Å).

Simulations were performed in LAMMPS. Structured domains were treated as rigid bodies and coupled to a Nosé–Hoover thermostat, whereas disordered regions were propagated using Langevin dynamics. Both used a relaxation time of 5 ps and a 10-fs integration timestep. For direct-coexistence simulations, protein replicas were initially arranged in a slab with a cross-sectional area of approximately 20 × 20 nm², yielding an average protein density of approximately 0.1 g·cm⁻³. Production simulations were run for 2 μs.

Intermolecular contact maps were calculated from trajectories at 290 K. Residue pairs were considered in contact when their separation was ≤1.2σᵢⱼ, slightly above the interaction-potential minimum (∼1.122σᵢⱼ), thereby restricting the analysis to substantially interacting residue pairs (Tejedor et al. 2021; Pedraza et al. 2025). Contacts per residue were normalized to protein length; for heterotypic systems, normalization was performed using the length of the larger protein to enable consistent comparison across systems.

## Statistical Analysis

Statistical analyses were performed using non-parametric methods. The Mann–Whitney U test (Wilcoxon rank-sum test) was used for pairwise comparisons. For differential gene expression, LGR5-high and LGR5-low tumors were compared, with significance defined as p ≤ 0.001, fold change (FC) ≥1.5, and expression ≥500 RNA-seq counts in the LGR5-high group. For somatic mutation analyses, LGR5 expression was compared between mutant and wild-type tumors; analyses were restricted to alterations present in ≥5 tumors, with p ≤ 0.05 and FC ≥1.44 used for prioritization. For copy-number analyses, LGR5 expression was compared between tumors harboring amplifications or deletions and copy-number-neutral tumors. Analyses were restricted to events present in ≥15 tumors; p ≤ 0.05 was required, together with FC ≥1.44 for amplifications.

P values should be interpreted as nominal, and complementary FC and prevalence thresholds were used to prioritize associations with larger effect sizes rather than to control the false discovery rate. Sample size for each analysis corresponded to the number of evaluable tumors with the required molecular data. Statistical analyses were performed using custom R scripts.

## Results

### LGR5 is overexpressed in colorectal cancer across pan-cancer analysis

Pan-cancer analysis revealed a restricted pattern of LGR5 expression, with the highest levels in rectal (READ, 21.17 TPM) and colon adenocarcinoma (COAD, 14.41 TPM), followed by uterine endometrial carcinoma (UCEC, 13.89 TPM) and uterine carcinosarcoma (UCS, 9.25 TPM) (Figure 1A). Most other tumor types showed low or negligible expression, consistent with the tissue-restricted distribution of LGR5 (Barker et al. 2007). Tumor-to-normal enrichment was particularly pronounced in UCEC, READ, UCS, PAAD, COAD, and ESCA (>20-fold), with minimal expression in matched normal tissues (Figure 1B). Together, these data identify colorectal cancer among the tumor types with the strongest and most selective LGR5 expression.

**Figure 1.**
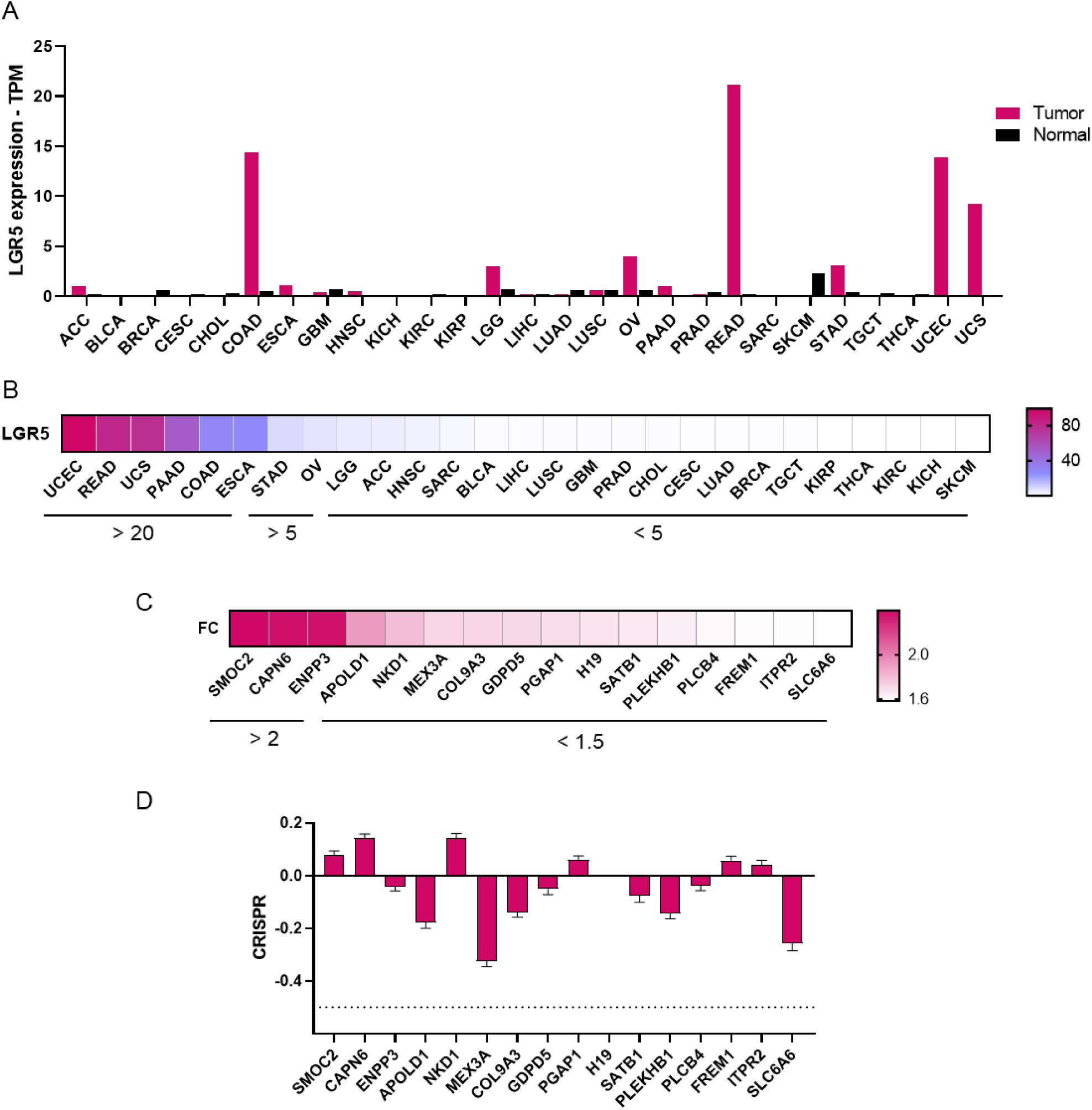
LGR5 expression across tumour types and characterisation of the LGR5-associated signature in colorectal adenocarcinoma. (A) Median LGR5 expression in transcripts per million across TCGA tumour cohorts and corresponding normal tissue; highest tumour expression is observed in READ, COAD, UCEC and UCS. (B) Heat map of the LGR5 tumour-to-normal expression ratio across the same cohorts, ordered by decreasing ratio, with bars indicating ratios above 20, above 5, and below 5. (C) Heat map depicting fold change between LGR5-high and LGR5-low COAD tumours for the genes of the LGR5-associated signature, ordered by decreasing fold change. (D) CRISPR dependency score (Chronos) for the same genes across colorectal cancer cell lines that overexpress LGR5. The dotted line indicates −0.5, below which a gene is conventionally considered essential. No signature gene crosses this threshold, indicating the absence of a selective genetic dependency.

### LGR5-high tumors displayed a coordinated transcriptional program linked to stemness and Wnt signaling

Comparison of LGR5-high (≥500 RNA-seq counts) and LGR5-low COAD tumors identified a distinct transcriptional program (p ≤ 0.001; FC ≥1.5). SMOC2, CAPN6, and ENPP3 showed the strongest enrichment (FC >2.3), while APOLD1, NKD1, MEX3A, COL9A3, and GDPD5 were consistently upregulated (FC >1.7) (Figure 1C). Additional genes, including PGAP1, SATB1, PLEKHB1, and ITPR2, further defined this expression state. Despite their coordinated upregulation, genome-wide CRISPR screening revealed no selective dependency on these genes in LGR5-high colorectal cancer cells (Figure 1D), indicating that the LGR5-high transcriptional program does not translate into an intrinsic genetic vulnerability.

### LGR5-high tumors are enriched for stemness, Wnt signaling, and surfaceome-associated programs

Functional annotation of the LGR5-high signature revealed coordinated enrichment of Wnt signaling, stemness, and tumor–microenvironment programs. NKD1, a negative Wnt regulator, was increased (FC = 1.8; p = 7.90 × 10⁻⁷), consistent with activation of Wnt feedback (Yan et al. 2001; Zeng et al. 2000), while the stemness-associated genes MEX3A and H19 and the matrix components COL9A3 and FREM1 supported enhanced cellular plasticity and stromal remodeling (Figure 1C) (Barriga et al. 2017; Ren et al. 2018; Naba et al. 2012). Surfaceome analysis further identified ENPP3 (FC = 2.4), GDPD5 (FC = 1.7), and SLC6A6 (FC = 1.6) as membrane-accessible components of the LGR5-high signature (Figure 1C) (Gorelik et al. 2018; Matas-Rico et al. 2016; Uchida et al. 1992), revealing candidate surface vulnerabilities within this transcriptional state.

### Somatic mutations and CNV associated with LGR5-high tumors define a distinct genomic context

Mutation analysis (n = 396) identified genomic alterations associated with elevated LGR5 expression (≥5 cases; p ≤ 0.05; FC ≥1.44). APC mutations were most prevalent (n = 291) and associated with increased LGR5 expression (FC = 1.62; p = 8.08 × 10⁻⁵), consistent with Wnt pathway activation (Figure S1A). Mutations in PCDHA3 and TGIF1 showed similar associations, whereas less frequent alterations in SLC44A2, KRT12, LAP3, SLC5A11, KCNMB1, and particularly SSX3 displayed larger effect sizes (FC up to 3.01). Despite their association with the LGR5-high state, CRISPR screening revealed no selective dependency on these genes in LGR5-high colorectal cancer cells (Figure S1B).

CNV analysis identified amplifications associated with elevated LGR5 expression, including IGF2 and INS-IGF2 (FC ∼1.69; p ≈ 0.024), together with KCNMB1, NFATC2, ATP9A, and recurrent BPIF family members (Figure S1C). Conversely, deletions of LAPTM5, MATN1, and SDC3 were associated with increased LGR5 expression (FC ∼1.68; p ≈ 0.03) (Figure S1E). These findings define a distinct copy-number landscape associated with the LGR5-high transcriptional state. However, genome-wide CRISPR screening revealed no selective dependency on these genes in LGR5-high colorectal cancer cells (Figure S1D-F).

### Comparative assessment of surface-accessible candidates for antibody-based targeting

To prioritize surface proteins enriched in LGR5-high tumors, we evaluated ENPP3 (CD203c), GDPD5 (GDE2), and SLC6A6 (TauT) using established determinants of ADC target suitability: tumor-to-normal selectivity, extracellular epitope accessibility, internalization and shedding behavior, and translational precedent (Ocana et al. 2026) (Table 1). Membrane-embedded structural models were used to contextualize extracellular accessibility according to protein topology (Figure 2, extracellular, transmembrane, and cytoplasmic regions shown in blue, orange, and green, respectively).

**Table 1.** Comparative assessment of the three surface-accessible candidates as antibody–drug conjugate targets. Qualitative comparison of ENPP3 (CD203c), GDPD5 (GDE2) and SLC6A6 (TauT) across four criteria relevant to ADC development: tumour selectivity, structural accessibility of the extracellular region, internalisation and shedding behaviour, and translational readiness, together with an overall suitability call. ENPP3 combines restricted normal-tissue expression, a large accessible ectodomain with experimentally determined structures, documented antibody-mediated lysosomal internalisation, and existing clinical ADC and bispecific precedents. GDPD5 retains an accessible extracellular catalytic domain and constitutive endocytosis despite a six-transmembrane topology, but shows only moderate tumour enrichment and no antibody-targeting precedent. SLC6A6 combines broad normal-tissue expression with a twelve-transmembrane architecture that leaves a limited extracellular surface, and its internalisation and shedding behaviour are uncharacterised. The overall assessment is a qualitative integration of the four criteria by the authors and is not a computed score.

| Target | Tumor Selectivity | Structural Accessibility | Internalization / Dynamics | Translational Readiness | Overall |
| --- | --- | --- | --- | --- | --- |
| ENPP3 (CD203c) | High tumor selectivity with restricted tissue expression | Large extracellular domain, single-pass membrane protein, experimental structures available | Rapid antibody-mediated lysosomal internalization; no major shedding concerns | Clinical ADC and bispecific precedents available | High |
| GDPD5 (GDE2) | Moderate tumor enrichment but broad normal tissue expression | Extracellular catalytic domain accessible despite six-transmembrane topology | Constitutive endocytosis described, mainly in neuronal systems | No ADC or antibody-targeting precedent | Medium |
| SLC6A6 (TauT) | Tumor overexpression offset by widespread expression in normal tissues | Poor antibody accessibility due to 12-transmembrane architecture and limited extracellular regions | Internalization and shedding properties largely unknown | No biologic or ADC development precedent | Low |

**Figure 2.**
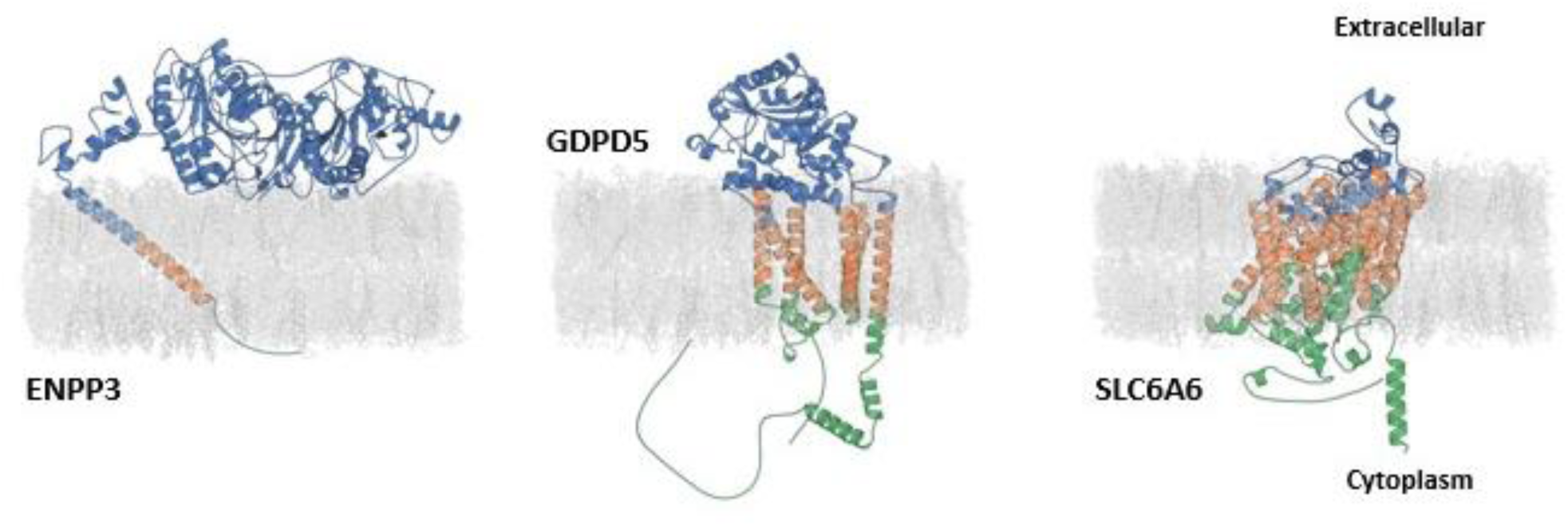
Comparative assessment of the three surface-accessible candidates as antibody–drug conjugate targets. Membrane-embedded structural models of each protein, shown in the same orientation with the extracellular compartment at the top and the cytoplasm at the bottom; the lipid bilayer is rendered in grey. Extracellular regions are coloured blue, transmembrane segments orange and intracellular regions green. Models were obtained as described in material and methods and positioned in the bilayer. For ENPP3, the orientation of the ectodomain relative to the membrane is not constrained by the single transmembrane helix and is shown schematically. The models illustrate the difference in accessible surface between the single-pass ENPP3, whose ectodomain lies entirely outside the bilayer, and the polytopic GDPD5 and SLC6A6, whose extracellular loops are short and closely opposed to the membrane.

Evaluation against established determinants of ADC target suitability clearly prioritized ENPP3. ENPP3 combines restricted normal-tissue expression (Figure 3A–B), a large and structurally validated extracellular domain providing broad epitope accessibility (Figure 2) (Gorelik et al. 2018), efficient antibody-mediated internalization and lysosomal trafficking, and limited ectodomain shedding (Doñate et al. 2016). Its tractability is further supported by clinical-stage ENPP3-directed ADC and bispecific programs (Doñate et al. 2016; Thompson et al. 2018; Kollmannsberger et al. 2021).

**Figure 3.**
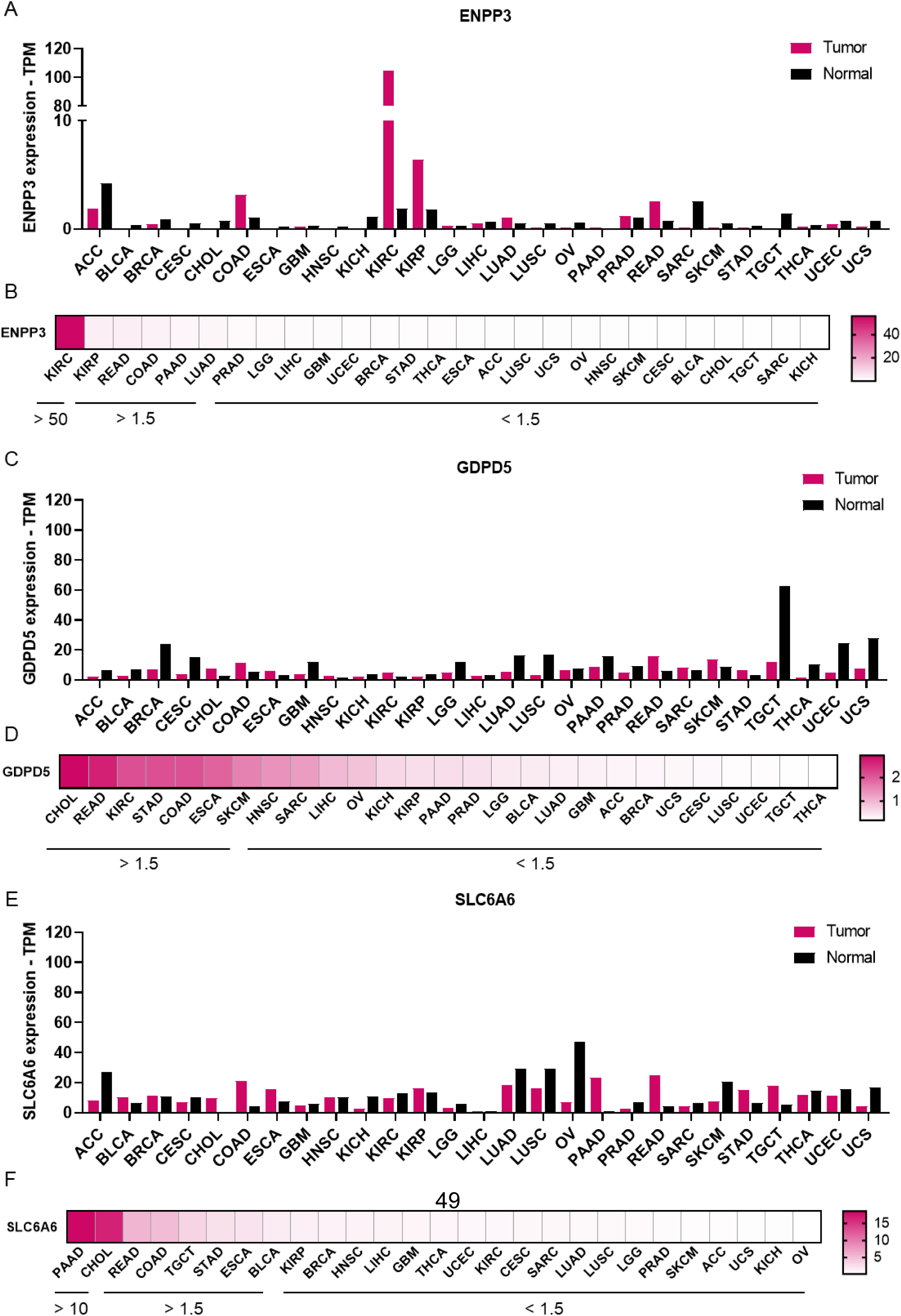
Pan-cancer tumour-versus-normal expression of the three surface-accessible candidates. Expression of ENPP3 (A, B), GDPD5 (C, D) and SLC6A6 (E, F) across TCGA tumour cohorts and matched normal tissue. (A, C, E) Median expression in transcripts per million (TPM) in tumour (magenta) and normal (black) tissue for each of the 27 TCGA cohorts, identified by their standard study abbreviations. The y-axis is fixed at 0–120 TPM across all three genes to allow direct comparison of absolute expression between candidates. (B, D, F) Heat map depicting fold change between tumor and non-transformed tissue, with cohorts ordered left to right by decreasing ratio; colour scales are gene-specific and are given to the right of each panel. Horizontal bars beneath each heat map group cohorts by ratio band: ENPP3 shows a ratio above 50 in KIRC and SLC6A6 (F) reach a maximum band of above 10, in PAAD and CHOL; the intermediate band (ratio > 1.5) and the non-enriched band (ratio < 1.5) are indicated in each case. ENPP3 displays the most restricted profile, with high absolute tumour expression confined to KIRC and near-absent expression across most normal tissues, whereas GDPD5 and SLC6A6 are broadly expressed in both tumour and normal tissue, with normal expression exceeding tumour expression in several cohorts (for example OV for SLC6A6 and TGCT for GDPD5).

GDPD5 showed intermediate tractability. Its extracellular catalytic domain is accessible despite its six-transmembrane topology (Figure 2), but broader normal-tissue expression limits tumor selectivity (Figure 3C–D) (Lang et al. 2008). Although constitutive internalization has been reported (Salgado-Polo et al. 2020), tumor-cell trafficking remains poorly defined, and no ADC or antibody-development precedent exists.

SLC6A6 showed the least favorable profile. Broad normal-tissue expression (Figure 3E–F), together with a 12-transmembrane topology exposing only short extracellular loops (Figure 2) (Du et al. 2025; Stary and Bajda 2024), limits both selectivity and epitope accessibility. Moreover, internalization and shedding remain undefined, with no reported ADC precedent. Thus, established ADC tractability parameters nominate ENPP3 as the strongest near-term candidate, while GDPD5 requires further validation and SLC6A6 lacks key features required for antibody-based targeting.

### Structure-based druggability assessment of candidate targets

The remaining upregulated targets—SMOC2, CAPN6, APOLD1, NKD1, SATB1, MEX3A, PGAP1, COL9A3, PLEKHB1, PLCB4, ITPR2, and FREM1—were evaluated for structural druggability using three complementary pocket-detection approaches. P2Rank, implemented through PrankWeb (Polák et al. 2025; Krivák and Hoksza 2018), predicts ligand-binding cavities using structural and evolutionary features; pockets with P2Rank scores ≥10.0 and probabilities ≥0.5 are reported in Table S1. DeepPocket combines Fpocket cavity detection with 3D convolutional neural-network re-ranking (Aggarwal et al. 2022; Le Guilloux, Schmidtke, and Tuffery 2009; Schmidtke and Barril 2010); pockets meeting CNNconfidence ≥0.5 and druggability ≥0.5 are summarized in Table S2. DoGSiteScorer independently evaluates pocket geometry and physicochemical properties (Volkamer et al. 2012), with cavities meeting SimpleScore and DrugScore ≥0.6 reported in Table S3. Integration across these complementary methods prioritized PLCB4, SATB1, PLEKHB1, SMOC2, and PGAP1 for further structural analysis. Pocket metrics were then integrated with functional annotation, structural context, and cancer relevance to derive an overall tractability assessment (Table 2).

**Table 2.**
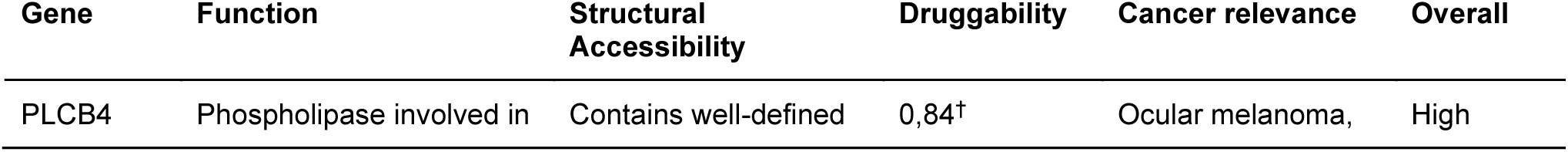
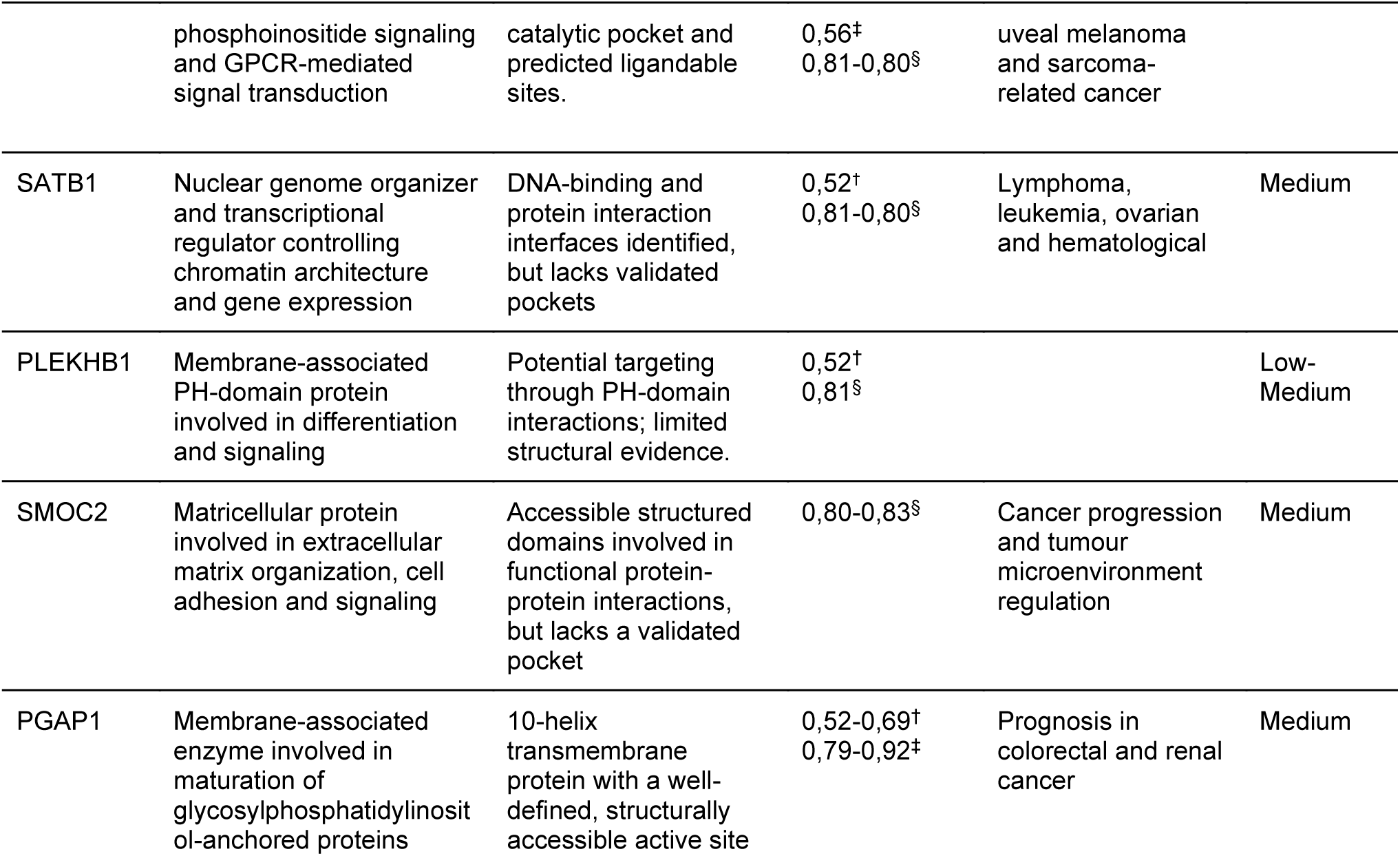
Intracellular candidates upregulated in LGR5-high tumours and computational assessment of their binding-site druggability. (A) Summary of biological function, structurally and functionally relevant regions supported by experimental evidence, predicted pocket druggability, and reported cancer relevance for PLCB4, SATB1, PLEKHB1, SMOC2 and PGAP1. Druggability scores were obtained with PrankWeb (†), DeepPocket (‡) and DoGSiteScorer (§); for proteins with multiple predicted binding pockets, values indicate the range observed across sites. Scores from the three methods are not on a common scale and should be compared within, not across, methods. Cancer relevance is drawn from GeneCards (Stelzer et al. 2016) and the scientific literature for each target. The overall assessment is a qualitative integration of the preceding columns by the authors and is not a computed score.

PLCB4 showed the strongest cross-method support, with all three approaches identifying its known active site and two additional putative cavities (Figure 4). PLCB4 regulates phosphoinositide and GPCR signaling and has been implicated in uveal melanoma and sarcoma, although its role in colorectal cancer remains undefined (Phan et al. 2021; Liu et al. 2025). In SATB1, multiple approaches recovered the known DNA-binding site and identified additional putative cavities (Figure 4). This chromatin organizer has been implicated across hematologic and solid malignancies (Cai, Han, and Kohwi-Shigematsu 2003; Dickinson, Dickinson, and Kohwi-Shigematsu 1997; Bai et al. 2025). PLEKHB1, a membrane-associated PH-domain protein involved in differentiation and signaling, contained a novel pocket independently supported by two approaches (Figure 4) (Xu et al. 1999).

**Figure 4.**
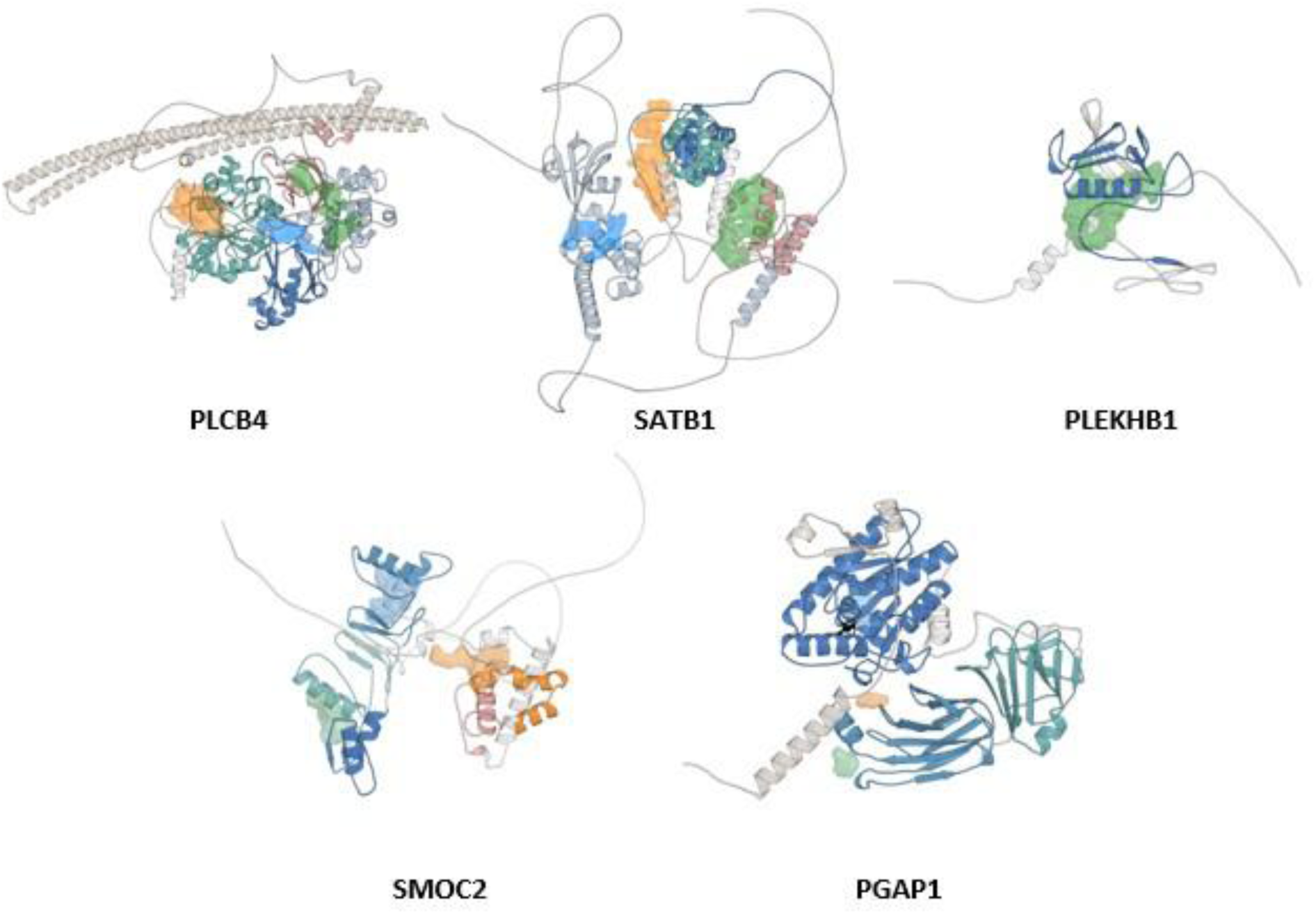
Intracellular candidates upregulated in LGR5-high tumours and computational assessment of their binding-site druggability. Representative structural models of PLCB4, SATB1, PLEKHB1 (top row, left to right) and SMOC2, PGAP1 (bottom row, left to right), obtained from AF-Q15147-F1-, AF-Q01826-F1, AF-Q9UF11-F1, AF-Q9H3U7-F1, and AF-Q75T13-F1, respectively, and rendered in PyMOL (Schrödinger, LLC, n.d.). Structured domains are coloured individually, with intrinsically disordered and low-confidence regions shown in grey.

SMOC2 contained two predicted pockets, including the reported Ca²⁺-binding site and a putative novel cavity (Figure 4). This matricellular protein regulates extracellular matrix organization, adhesion, and tumor–microenvironment signaling (Maier, Paulsson, and Hartmann 2008; Shvab et al. 2016; Chiodoni, Colombo, and Sangaletti 2010). PGAP1 similarly contained its catalytic pocket and an additional predicted cavity (Figure 4); however, the latter localized to a transmembrane region, reducing its tractability despite cross-method support (Hong et al. 2024; Tanaka et al. 2004). Remaining targets were deprioritized because of transmembrane pocket localization (APOLD1), poorly defined cavities (MEX3A), limited cross-method support (COL9A3), or incomplete structural information (ITPR2).

Among the five candidates, SATB1 showed the strongest tumor-associated expression, driven by markedly elevated levels in LGG (∼3,000 TPM) with substantially lower expression in matched normal tissue (Figure S2C–D). PLCB4 showed a more selective pattern, with tumor enrichment in COAD and READ (Figure S2A–B). PLEKHB1 displayed heterogeneous expression, including strong enrichment in LGG but higher normal-tissue expression in selected contexts such as GBM (Figure S2E–F). SMOC2 generally showed limited tumor selectivity, with normal tissues frequently expressing comparable or higher levels despite enrichment in CHOL and other tumors (Figure S2G–H). PGAP1 showed the weakest differential, with low overall expression and only modest tumor enrichment (Figure S2I–J). Thus, SATB1 and PLCB4 displayed the most favorable tumor-associated expression profiles, whereas SMOC2 and PGAP1 showed limited tumor selectivity.

### Condensate propensity extends druggability assessment beyond surface-accessible targets and structure-based pocket druggability

Beyond classical pocket-based druggability, several of the genes identified in the LGR5-high signature lacked a tractable small-molecule binding site, or displayed only a moderate, function-specific cavity, limiting their accessibility to conventional structure-based drug design. For such targets, modulating their capacity to form biomolecular condensates or targeting their intrinsically disordered regions offers a complementary therapeutic strategy (Mitrea et al. 2022), particularly relevant for genes implicated in Wnt pathway regulation, given that the Wnt signalling machinery itself relies on phase-separation-driven assemblies (Nong et al. 2021; Kang et al. 2022). We therefore assessed the homotypic phase-separation propensity of genes from the LGR5-high signature as an alternative route to identify pharmacologically actionable vulnerabilities.

Saturation concentration (c_sat_) predictions restricted to the IDR fraction of each protein passing the disorder threshold described in the methods section did not identify any candidate with a c_sat_ value compatible with homotypic phase separation (Figure 5A). Based on these predictions together with the net charge of each protein (Figure 5B), three candidates were selected for Direct Coexistence (DC) molecular dynamics simulations: SATB1 and MEX3A, both previously reported to form condensates in presence of other scaffold biomolecules (Madhok et al. 2025; Chen et al. 2024), and NKD1, prioritized on the basis of its predicted c_sat_ and its high net positive charge, a feature associated with the formation of protein–RNA complex coacervates (Espejo et al. 2025; Banerjee et al. 2017; Maharana et al. 2018).

**Figure 5.**
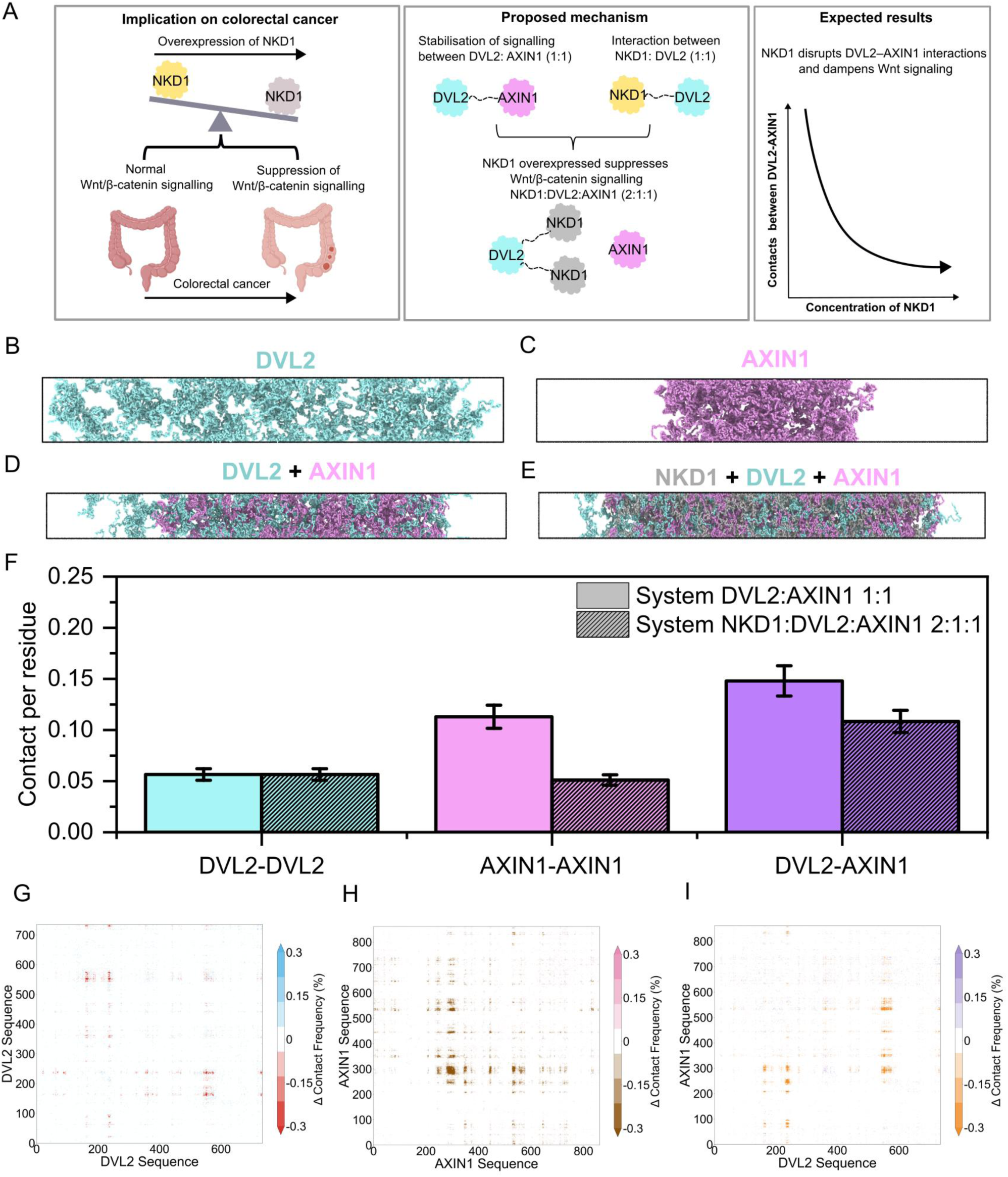
NKD1 partitions into DVL2 condensates and weakens DVL2–AXIN1 association. (A) Schematic of the proposed mechanism: NKD1 overexpression suppresses Wnt/β-catenin signalling in colorectal cancer by interacting with AXIN1 (1:1) and disrupting stabilising DVL2– AXIN1 (1:1) contacts, dampening Wnt signalling output as NKD1 concentration increases (NKD1:DVL2: AXIN1, 2:1:1). (B-C) Representative slab snapshots from direct-coexistence simulations of DVL2 (teal) and AXIN1 (magenta) alone. (D) DVL2 + AXIN1 binary system (1:1). (E) Ternary NKD1 (grey) + DVL2 (teal) + AXIN1 (magenta) system (2:1:1). (F) Intermolecular contacts per residue for DVL2-DVL2, AXIN1-AXIN1, and DVL2-AXIN1 pairs in the binary (DVL2:AXIN1 1:1, solid) versus ternary (NKD1:DVL2:AXIN1 2:1:1, hatched) systems. (G-I) Residue-resolved difference contact maps (Δ contact frequency, ternary minus binary system) for DVL2-DVL2 (G), AXIN1-AXIN1 (H), and DVL2-AXIN1 (I) interfaces. Error bars in (F) represent standard error of the mean.

### NKD1 undergoes homotypic phase separation and forms RNA-stabilised condensates

Simulations of the isolated proteins using our coarse-grained Mpipi-Recharged model (R. Tejedor et al. 2025) (described in the methods section) showed that NKD1 undergoes homotypic phase separation at 290 K and physiological salt concentration, with an average of ∼30% intermolecular contacts per residue, whereas SATB1 and MEX3A failed to form stable pure protein condensates, displaying only ∼5% intermolecular contacts per residue (Figure 5C–D). Addition of single stranded polyA RNAs using the compatible Mpipi-Recharged RNA nucleotide-resolution model (Tejedor et al. 2026), titrated to the isoelectric point of the NKD1 mixture and matched in amount across the remaining two systems for consistency, stabilized the NKD1 condensate through extensive protein–RNA contacts. In contrast, although SATB1 and MEX3A established protein–RNA contacts of comparable magnitude (∼5%), this was insufficient to drive phase separation in either system (Figure 5E– F). Residue-residue and residue-RNA contact maps for all systems are provided in Figure S3, highlighting the interaction patterns underlying these distinct condensation propensities.

### NKD1 partitions into DVL2 condensates and weakens DVL2–AXIN1 association

NKD1 is a Wnt-inducible negative feedback regulator that antagonises Wnt/β-catenin signalling by binding Dishevelled (DVL) family scaffold proteins through its EFX domain (Wharton et al. 2001; Zhang et al. 2007). NKD1 mutations that disrupt this interaction have been identified in mismatch repair-deficient colorectal tumours, where they result in increased Wnt/β-catenin signalling (Guo et al. 2009), underscoring the functional importance of the NKD1–DVL interaction in limiting Wnt pathway activity. In binary DC simulations, NKD1 engaged both DVL2 and AXIN1 directly, with comparable NKD1–DVL2 and NKD1–AXIN1 contacts per residue (Figure S4A–B), indicating that NKD1 has the intrinsic capacity to partition into condensates formed by either scaffold. The Wnt/β-catenin pathway is itself organised by condensate formation at two opposing nodes: in the Wnt OFF state, the β-catenin destruction complex assembles through APC-promoted, IDR-driven LLPS of AXIN1, concentrating the kinases GSK3β and CK1α to drive β-catenin degradation (Nong et al. 2021). Upon Wnt stimulation, DVL2 itself undergoes IDR-mediated condensation to nucleate the receptor signalosome, sequestering AXIN1 away from the destruction complex and thereby allowing β-catenin to accumulate (Kang et al. 2022). Given that NKD1 directly binds DVL2, the principal scaffold of this condensate system, we asked whether NKD1 partitions into DVL2 condensates and whether this affects DVL2–AXIN1 association — a relationship that has not been previously tested (Figure 6A). To address this, we performed DC simulations of NKD1, DVL2, and AXIN1, evaluating each protein alone and in combination.

**Figure 6.**
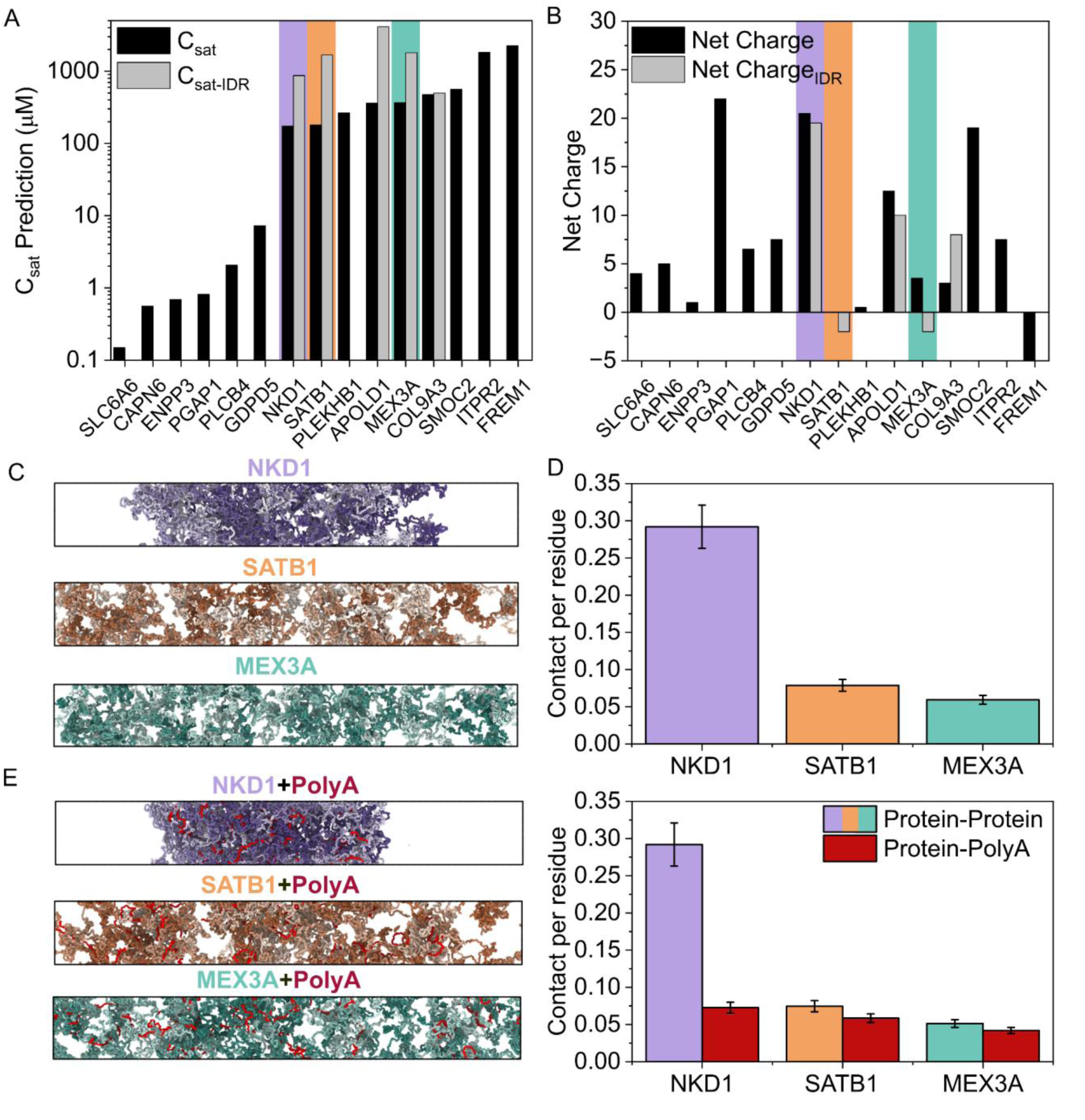
NKD1 undergoes homotypic phase separation and forms RNA-stabilised condensates, unlike SATB1 and MEX3A. (A) Predicted saturation concentration (c_sat_) for the full-length sequence (black) and the IDR-only sequence (grey) of each candidate, ranked by full-length c_sat_. Bars highlighted in purple, orange, and teal correspond to NKD1, SATB1, and MEX3A, respectively. (B) Net charge of the full-length sequence (black) and the IDR (grey) for each protein, with NKD1, SATB1, and MEX3A highlighted as in (A). (C) Representative slab snapshots from direct-coexistence simulations of NKD1 (purple), SATB1 (orange), and MEX3A (teal) alone. (D) Intermolecular protein-protein contacts per residue for NKD1, SATB1, and MEX3A, averaged over the production trajectory. (E) Representative slab snapshots of NKD1, SATB1, and MEX3A in the presence of polyA RNA (red), and corresponding protein-protein (coloured) versus protein-RNA (red) contacts per residue. Error bars in (D) and (E) represent standard error of the mean.

Our simulations show that AXIN1, but not DVL2 (Figure 6B-C), phase separates when simulated individually. Upon mixing, the two proteins co-condense into a mixed DVL2–AXIN1 assembly at 1:1 stoichiometry (Figure 6D). Addition of NKD1 into the binary mixture (at a 2:1:1 NKD1:DVL2:AXIN1 stoichiometry) produced a markedly different condensate architecture (Figure 6E): NKD1 co-localised extensively with both DVL2 and AXIN1, with NKD1–DVL2 contacts becoming more prominent than NKD1–AXIN1 contacts in the ternary system (Figure S4C), consistent with a direct client-like interaction with both scaffolds. This recruitment was accompanied by a pronounced decrease in AXIN1–AXIN1 self-contacts (∼50%) and a more modest decrease in DVL2–AXIN1 contacts (∼25%) relative to the binary DVL2:AXIN1 system, while DVL2–DVL2 self-contacts remained essentially unchanged (Figure 6F). Residue-resolved contact map differences between the distinct condensate mixtures showed that NKD1 recruitment reduces contact frequency across extended regions of the AXIN1 sequence in the AXIN1–AXIN1 and DVL2–AXIN1 interfaces, while leaving the DVL2–DVL2 interface largely unaffected (Figure 6G–I). Residue-resolved intermolecular contact maps for all simulated systems are presented in Figure S5, spanning DVL2-only, AXIN1-only, binary DVL2:AXIN1 (1:1), binary NKD1:DVL2 (1:1), binary NKD1:AXIN1 (1:1), and ternary NKD1:DVL2:AXIN1 (2:1:1) systems. Figure S4 summarises contacts per residue across all homotypic and heterotypic pairs (NKD1–NKD1, DVL2–DVL2, AXIN1–AXIN1, DVL2–AXIN1, NKD1–DVL2, and NKD1–AXIN1), comparing individual protein condensate simulations against their binary and ternary counterparts to highlight stoichiometry-dependent effects on interaction patterns. Collectively, our results indicate that NKD1 is recruited into the DVL2–AXIN1 condensate through direct interactions with both scaffolds and, at sufficient concentration, preferentially disrupts AXIN1 self-association and DVL2– AXIN1 contacts while leaving DVL2 self-association largely intact, providing a biophysical mechanism by which NKD1 overexpression could dampen Wnt signalling in LGR5-high colorectal tumours.

## Discussion

Here, we define the multi-omic landscape of LGR5-high colorectal cancer, revealing a coordinated molecular state characterized by Wnt pathway activation, intestinal stemness, and extracellular matrix remodeling (Barker et al. 2007; Schepers et al. 2012; Muzny et al. 2012). By integrating transcriptional, mutational, and copy-number features with three complementary dimensions of pharmacological tractability, we translate this molecular state into distinct therapeutic vulnerabilities addressable through orthogonal modalities.

The LGR5-high transcriptional program is consistent with a Wnt-active, stem-like cellular state, in line with the established role of LGR5 as a canonical Wnt target and intestinal stem cell marker (Barker et al. 2007; de Lau et al. 2011). Enrichment of SMOC2, a marker of LGR5-positive crypt-base stem cells (Muñoz et al. 2012), and MEX3A, which defines a slow-cycling population capable of tumor reinitiation after chemotherapy (Pereira, Billaud, and Almeida 2017), reinforces this stem-like phenotype and suggests a potential link to therapeutic persistence. Increased NKD1, a negative-feedback regulator induced by Wnt/β-catenin signaling, further supports sustained pathway activity, while enrichment of the oncofetal lncRNA H19 (Raveh et al. 2015) points to increased cellular plasticity. Together, these features define LGR5-high colorectal cancer as a coordinated Wnt-driven, stem-like transcriptional state rather than the consequence of LGR5 upregulation alone.

Notably, none of the coordinately upregulated genes emerged as selective CRISPR dependencies in LGR5-high colorectal cancer cells. Such disconnection between transcriptional enrichment and genetic essentiality does not preclude therapeutic relevance, but instead distinguishes dependency from tractability. ADC targets, for example, require tumor-selective expression, accessibility, and internalization rather than oncogenic dependency, as illustrated by the activity of trastuzumab deruxtecan in HER2-low tumors (Modi et al. 2022). Similarly, microenvironmental proteins such as SMOC2, COL9A3, and FREM1 are poorly captured by monoculture viability screens, while loss of negative-feedback regulators such as NKD1 may enhance rather than suppress pathway output. Thus, the absence of selective dependencies argues against conventional oncogene-targeting strategies but supports a modality-matched framework that exploits surface accessibility, ligandability, or biophysical behavior.

The enrichment of surface-accessible molecules in LGR5-high tumours represents a therapeutically compelling finding, but our comparative assessment indicates that surfaceome membership alone is a poor guide to tractability. Of the three surface-annotated candidates evaluated, only ENPP3 satisfied all four criteria relevant to ADC development. ENPP3 is a type II transmembrane glycoprotein with established expression on basophils and mast cells and limited expression in normal colonic epithelium (Tsai and Takeda 2016). Its single-pass topology presents a large globular ectodomain well clear of the bilayer, and experimentally determined structures of the human ectodomain are available to support rational epitope selection (Gorelik et al. 2018). ENPP3 has been the subject of a clinical ADC programme in renal cell carcinoma, where AGS-16C3F, an anti-ENPP3 antibody conjugated to monomethyl auristatin F, established that the antigen undergoes efficient antibody-mediated internalisation and lysosomal delivery in patients (Doñate et al. 2016; Thompson et al. 2018). That programme did not meet its randomised phase 2 endpoint. However, the investigators explicitly excluded target expression as the cause, reporting a median ENPP3 H-score of 255 of 300 in the treated population, and attributed the outcome instead to the limited chemosensitivity of renal cell carcinoma and to a heterogeneous enrolled population (Kollmannsberger et al. 2021). More recently, ENPP3-directed bispecific T-cell engagers have shown objective responses in the same indication, providing independent evidence that the antigen supports clinically meaningful tumour cell killing. Taken together, this history supports ENPP3 as a validated delivery portal whose first clinical asset failed on payload class and indication rather than on antigen biology — a distinction directly relevant to colorectal cancer, which, unlike renal cell carcinoma, is sensitive to topoisomerase I inhibition.

By contrast, GDPD5 and SLC6A6 expose the limitations of annotation-based surface target selection. GDPD5 presents an accessible extracellular domain, but modest tumor enrichment, broad normal-tissue expression (Lang et al. 2008), and poorly characterized trafficking in epithelial cells constrain its ADC potential. SLC6A6 provides an even clearer example: despite robust transcriptional enrichment, its widespread normal-tissue expression and 12-transmembrane topology, which exposes only short extracellular loops, make it unfavorable for antibody-based targeting (Uhl and PauL R. 1992). Notably, its substrate-binding cavity scored highly in pocket-detection analyses, potentially redirecting SLC6A6 toward small-molecule intervention. Thus, surface annotation alone does not define therapeutic modality; topology, tumor selectivity, and target-specific biology determine whether a candidate is tractable and how it should be targeted.

Structure-based analysis stratified the intracellular candidates by ligandability. CAPN6, NKD1, COL9A3, APOLD1, and FREM1 lacked cavities compatible with conventional small-molecule targeting, exemplified by the extended, pocket-poor architecture of COL9A3 and the transmembrane localization of predicted APOLD1 cavities. MEX3A similarly lacked well-defined druggable pockets. A second tier, including SATB1 and PLEKHB1, contained potentially tractable but higher-risk sites. The dominant SATB1 pocket maps to its DNA-binding interface, suggesting that pharmacological intervention would require disruption of macromolecular interactions, although SATB1 depletion suppresses colorectal tumorigenesis (Mir et al. 2016). PLEKHB1 contained a putative PH-domain cavity supported by two independent methods, consistent with the emerging druggability of PH domains (Powis et al. 2023), but lacking experimental validation. Thus, these targets are not intrinsically undruggable, but their reliance on interface-directed or unvalidated binding sites substantially increases discovery risk.

A third tier contained well-defined ligandable sites, with PLCB4 emerging as the strongest small-molecule candidate. PLCB4 ranked highest across all three pocket-detection methods, combining a defined catalytic site with additional predicted cavities. Its tractability is supported by oncogenic Gαq/11–PLCB4 signaling in melanoma (Phan et al. 2021), although achieving selectivity across closely related PLCβ isoforms remains a medicinal chemistry challenge (Lyon and Tesmer 2013) and its therapeutic relevance in colorectal cancer remains to be established. SMOC2 and PGAP1 were less compelling: SMOC2 contained a functionally supported Ca²⁺-binding cavity but showed unfavorable normal-tissue expression, whereas PGAP1 harbored a defined catalytic region but limited target precedent and an additional cavity embedded within its transmembrane architecture (Yang et al. 2016). Notably, several predicted pockets were absent from UniProt annotation (The UniProt Consortium 2025), highlighting the ability of structure-based approaches to uncover previously unannotated ligandable sites, although incomplete structural coverage, as observed for ITPR2, requires cautious interpretation.

The enrichment of APC mutations in LGR5-high tumors is consistent with canonical Wnt pathway biology. APC loss, present in ∼70–80% of sporadic colorectal cancers (Clevers and Nusse 2012), disrupts the β-catenin destruction complex, promoting nuclear β-catenin accumulation and transcription of Wnt targets including LGR5 (Barker et al. 2007; Fearon 2011). Accordingly, APC-mutant tumors showed significantly higher LGR5 expression (FC ∼1.62), supporting APC loss as a major genetic determinant of the LGR5-high state. Less frequent alterations, including SSX3, LAP3, and SLC44A2, were associated with larger LGR5 expression differences but occurred in small patient subsets and required validation. Thus, distinct genomic alterations may converge on a common Wnt-active, stem-like transcriptional state (Guinney et al. 2015).

The copy-number landscape provides an additional layer of therapeutic stratification. IGF2, which signals through IGF1R and insulin receptor pathways to activate PI3K–AKT and MAPK signaling (Pollak 2012), was amplified and upregulated in LGR5-high tumors, consistent with its established deregulation in colorectal cancer (Cui et al. 2003). This co-occurrence raises the possibility of an IGF2–IGF1R vulnerability within a molecularly defined subset, despite the limited efficacy of IGF1R-directed therapies in unselected colorectal cancer. Conversely, deletions involving LAPTM5, SDC3, and MATN1 were associated with the LGR5-high state. Although their functional significance remains unresolved, loss of regulators such as SDC3, which modulates growth-factor signaling and adhesion (Bernfield et al. 1999), may contribute to the signaling and cellular plasticity characteristic of this phenotype.

Beyond its transcriptional and genomic context, NKD1 emerged as a biophysically distinct component of the LGR5-high signature and illustrates the need for a third axis of therapeutic tractability. NKD1 is intracellular and lacks a conventional ligandable cavity, placing it beyond antibody-based and pocket-directed approaches. Although IDR-based C_sat_ predictions did not identify intrinsic condensation propensity, Direct Coexistence simulations revealed that full-length NKD1, unlike SATB1 and MEX3A, forms condensates that are further stabilized by RNA. This divergence highlights the potential limitations of IDR-restricted predictors in capturing condensation driven by full-length sequence context and heterotypic interactions (Espejo et al. 2025; Sanchez-Burgos et al. 2025).

This behavior is particularly relevant to Wnt signaling, as NKD1 directly binds DVL2 (Wharton et al. 2001; Guo et al. 2009), which organizes the receptor signalosome through phase separation (Kang et al. 2022). Ternary simulations showed that NKD1 partitions into DVL2 condensates and, with increasing stoichiometry, shifts DVL2 interactions away from AXIN1 and toward NKD1-associated assemblies. These findings suggest a biophysical mechanism for NKD1-mediated negative feedback in which condensation modulates DVL2 availability for productive Wnt signalosome assembly, complementing its established transcriptional feedback function (de Lau et al. 2011; Kang et al. 2022; Guo et al. 2009). More broadly, NKD1 illustrates how condensate biology may expose therapeutic opportunities among proteins inaccessible to conventional modalities (Mitrea et al. 2022). However, condensate-directed pharmacology remains early in development, and experimental validation will be required to establish both the proposed mechanism and its therapeutic tractability.

The principal advance of this study is the integration of multi-omic profiling with modality-matched therapeutic tractability. Rather than treating the LGR5-high molecular signature as a descriptive endpoint, we use it as the starting point for systematic prioritization across three orthogonal dimensions: surface accessibility, structural ligandability, and condensate phase behavior. This framework resolves distinct therapeutic opportunities that would not emerge from genomic or transcriptomic profiling alone, nominating ENPP3 for antibody-based targeting, PLCB4 for small-molecule intervention, and NKD1 for condensate-directed modulation. In the latter case, molecular simulations further uncover a biophysical mechanism whereby NKD1 condensation remodels the DVL2–AXIN1 interaction network, reducing AXIN1 self-association and DVL2–AXIN1 contacts and thereby potentially modulating Wnt signaling. Thus, integrating molecular state with target-specific biophysical properties provides a path from tumor profiling to modality-matched therapeutic hypotheses.

This study has several limitations. Bulk RNA-seq cannot resolve intratumoral heterogeneity, and the LGR5-high state may therefore reflect both cellular composition and tumor-intrinsic transcriptional programs; single-cell validation will be important to distinguish these contributions. The empirically defined LGR5 threshold (≥500 RNA-seq counts) may influence subgroup composition and should be validated in independent cohorts using continuous expression measures. Moreover, differential expression, mutation, and CNV analyses were not corrected for multiple testing, and despite complementary effect-size thresholds, false-positive associations cannot be excluded. Finally, therapeutic tractability was inferred computationally from structural models, curated annotations, and coarse-grained simulations rather than experimentally established. The nominated targets should therefore be considered prioritized therapeutic hypotheses requiring orthogonal validation, including protein expression and internalization for ENPP3, biochemical and selectivity profiling for PLCB4, and cellular validation of NKD1 condensate behavior and its impact on Wnt signaling.

### Conclusions

These findings define LGR5-high colorectal cancer as a distinct Wnt-active, stem-like molecular state shaped by APC loss and additional genomic alterations, yet lacking selective genetic dependencies. Integrating this signature with three orthogonal dimensions of therapeutic tractability nominates ENPP3 for antibody-based delivery, PLCB4 for small-molecule targeting, and NKD1 for condensate-directed modulation. More broadly, this framework moves beyond descriptive molecular profiling by matching tumor-associated vulnerabilities to the therapeutic modality best suited to their biological and biophysical properties. Experimental and spatial validation of these predictions will be required, but this approach provides a generalizable strategy for converting molecular tumor states into modality-matched therapeutic hypotheses.

## Acknowledgements

Ethics approval and consent to participate

Not applicable. This study used publicly available datasets and established human cancer cell lines, together with in silico protein simulations.

## Consent for publication

Not applicable.

## Availability of data and materials

The datasets used and/or analysed during the current study are available from the corresponding author on reasonable request.

## Authors’ contributions

LPH analyzed and interpreted the data regarding LGR5 expression, genetic alterations, CRISPR dependencies, and candidate therapeutic targets, and was a major contributor to writing the manuscript. AF performed, analyzed, and interpreted the data regarding CD simulations and biocondensates, and was a contributor to writing the manuscript. CP performed, analyzed, and interpreted the in silico data regarding the druggability of protein targets, and was a contributor to writing the manuscript. ART interpreted the data regarding CD simulations and biocondensates. LAC, BD, JAA, VA, CNJ, CAM, VM, EC, BG, JRE and AO supervised the study and were contributors to writing the manuscript. All authors read and approved the final manuscript.

## Appendices

Not applicable.

