## Supplementary Material for "LGR5-high colorectal cancer defines a Wnt-active stem-like state with multimodal therapeutic vulnerabilities"

**Affiliations**

| Gene | Structure | Pocket | Score | Probability | #Res | AvgCons | AvgAFscore |
|---|---|---|---|---|---|---|---|
| ENPP3 | AF-O14638-F1 | pocket1 | 20,7 | 0,80 | 26 | 1,27 | 98,4 |
| SATB1 | AF-Q01826-F1 | pocket1 | 10,3 | 0,53 | 18 | - | 88,3 |
| PGAP1 | AF-Q75T13-F1 | pocket1 | 15,0 | 0,69 | 18 | 0,00 | 85,5 |
| PGAP1 | AF-Q75T13-F1 | pocket2 | 14,6 | 0,68 | 16 | 3,08 | 96,2 |
| PGAP1 | AF-Q75T13-F1 | pocket3 | 11,9 | 0,59 | 15 | 0,00 | 86,9 |
| PGAP1 | AF-Q75T13-F1 | pocket4 | 10,0 | 0,52 | 10 | 2,90 | 94,2 |
| GDPD5 | AF-Q8WTR4-F1 | pocket1 | 28,4 | 0,88 | 28 | 1,21 | 94,7 |
| SLC6A6 | AF-P31641-F1 | pocket1 | 33,5 | 0,91 | 44 | - | 88,7 |
| PLEKHB1 | AF-Q9UF11-F1 | pocket1 | 10,2 | 0,52 | 14 | 1,95 | 73,0 |
| PLCB4 | AF-Q15147-F1 | pocket1 | 24,5 | 0,84 | 34 | - | 82,7 |

**Table S1.** Binding sites identified by PRANK with a score ≥ 10 and a probability ≥ 0.5. Descriptors: **score**, score assigned by the PRANK model; **probability**, estimated probability that the predicted site corresponds to a ligand-binding pocket; **#Res**, number of residues defining the binding site; **avgCons**, average evolutionary conservation score of the residues forming the pocket; and **avgAFscore**, average AlphaFold confidence score of the residues forming the pocket.

| Gene | Structure | Pocket | CNN confidence | Druggability | Volume | SASA | Polarity | HPh | Alpha spheres |
|---|---|---|---|---|---|---|---|---|---|
| APOLD1 | AF-Q96LR9-F1 | 21 | 0,82 | 0,51 | 809,8 | 248,9 | 6 | 33,9 | 103 |
| APOLD1 | AF-Q96LR9-F1 | 22 | 0,69 | 0,56 | 390,1 | 162,3 | 3 | 41,6 | 58 |
| PGAP1 | AF-Q75T13-F1 | 20 | 0,92 | 0,92 | 635,2 | 207,6 | 6 | 47,8 | 80 |
| PGAP1 | AF-Q75T13-F1 | 14 | 0,90 | 0,86 | 962,8 | 295,2 | 11 | 42,6 | 125 |
| PGAP1 | AF-Q75T13-F1 | 26 | 0,80 | 0,79 | 332,7 | 93,9 | 1 | 93,6 | 36 |
| GDPD5 | AF-Q8WTR4-F1 | 25 | 0,53 | 0,72 | 354,7 | 117,0 | 3 | 40,8 | 44 |
| SLC6A6 | AF-P31641-F1 | 5 | 0,99 | 0,90 | 1529,3 | 419,4 | 21 | 40,2 | 198 |
| ITPR2 | AF-Q14571-2-F1 | 18 | 0,81 | 0,76 | 992,8 | 328,9 | 5 | 42,7 | 82 |
| PLCB4 | AF-Q15147-F1 | 98 | 0,51 | 0,56 | 704,9 | 207,9 | 8 | 25,5 | 55 |
| FREM1 | AF-Q5H8C1-F1 | 178 | 0,69 | 0,62 | 554,7 | 196,7 | 6 | 30,2 | 46 |

**Table S2.** Binding sites identified by DeepPocket with a **CNNconfidence** ≥ 0.5 and a **druggability** ≥ 0.5. Descriptors: **CNNconfidence**, confidence score assigned by the CNN used for pocket classification; **druggability**, predicted probability that the pocket is druggable; **volume**, geometric volume of the pocket (in $Å^3$); **SASA**, solvent-accessible surface area of the pocket (in $Å^2$); **polarity**, polarity score of residues lining the pocket; **HPh**, hydrophobicity score of the pocket; and **Alpha spheres**, number of alpha spheres used to describe and represent the pocket geometry.

| Gene | Structure | Pocket | SimpleScore | DrugScore | Volume | Surface | Depth | HPh | siteAtoms |
|---|---|---|---|---|---|---|---|---|---|
| SMOC1 | AF-Q9H3U7-F1 | P_0 | 0,71 | 0,80 | 1337,3 | 2109,9 | 29,5 | 0,58 | 200 |
| SMOC1 | AF-Q9H3U7-F1 | P_1 | 0,64 | 0,83 | 890,1 | 1372,9 | 25,0 | 0,54 | 174 |
| SMOC1 | AF-Q9H3U7-F1 | P_2 | 0,62 | 0,83 | 861,9 | 1417,3 | 22,9 | 0,54 | 142 |
| APOLD1 | AF-Q96LR9-F1 | P_0 | 0,70 | 0,81 | 2456,8 | 3579,0 | 24,7 | 0,54 | 370 |
| APOLD1 | AF-Q96LR9-F1 | P_1 | 0,70 | 0,81 | 2059,8 | 3200,8 | 35,1 | 0,54 | 345 |
| NKD1 | AF-Q969G9-F1 | P_0 | 0,71 | 0,80 | 2102,0 | 2678,2 | 20,1 | 0,58 | 241 |
| NKD1 | AF-Q969G9-F1 | P_1 | 0,70 | 0,80 | 1244,7 | 1861,2 | 25,3 | 0,55 | 179 |
| SATB1 | AF-Q01826-F1 | P_0 | 0,66 | 0,80 | 2760,0 | 2759,5 | 35,8 | 0,43 | 351 |
| SATB1 | AF-Q01826-F1 | P_1 | 0,70 | 0,81 | 2654,2 | 3139,9 | 23,0 | 0,54 | 341 |
| SATB1 | AF-Q01826-F1 | P_2 | 0,71 | 0,80 | 2239,1 | 2905,6 | 31,0 | 0,57 | 282 |
| SATB1 | AF-Q01826-F1 | P_3 | 0,68 | 0,80 | 1680,0 | 2202,0 | 24,1 | 0,49 | 246 |
| MEX3A | AF-A1L020-F1 | P_0 | 0,70 | 0,81 | 5308,0 | 6766,3 | 39,5 | 0,56 | 680 |
| GDPD5 | AF-Q8WTR4-F1 | P_0 | 0,64 | 0,81 | 1504,7 | 1779,6 | 35,9 | 0,39 | 353 |
| GDPD5 | AF-Q8WTR4-F1 | P_1 | 0,67 | 0,84 | 890,4 | 1062,9 | 22,2 | 0,62 | 138 |
| COL9A3 | AF-Q14050-F1 | P_0 | 0,74 | 0,81 | 2741,8 | 4077,0 | 47,1 | 0,65 | 278 |
| COL9A3 | AF-Q14050-F1 | P_1 | 0,73 | 0,80 | 1096,9 | 1884,0 | 28,4 | 0,62 | 113 |
| SLC6A6 | AF-P31641-F1 | P_0 | 0,66 | 0,80 | 1338,0 | 1939,2 | 25,2 | 0,45 | 244 |
| SLC6A6 | AF-P31641-F1 | P_1 | 0,66 | 0,82 | 1281,5 | 1213,1 | 27,4 | 0,43 | 287 |
| PLEKHB1 | AF-Q9UF11-F1 | P_0 | 0,68 | 0,81 | 2642,2 | 3629,7 | 30,4 | 0,49 | 426 |
| PLCB4 | AF-Q15147-F1 | P_1 | 0,66 | 0,80 | 1850,1 | 2042,6 | 26,0 | 0,44 | 299 |
| PLCB4 | AF-Q15147-F1 | P_2 | 0,64 | 0,81 | 1455,3 | 1837,8 | 30,0 | 0,38 | 347 |
| PLCB4 | AF-Q15147-F1 | P_3 | 0,60 | 0,80 | 1318,5 | 1285,0 | 25,7 | 0,30 | 312 |
| PGAP1 | AF-Q75T13-F1 | P_0 | 0,60 | 0,81 | 1154,2 | 1169,3 | 23,8 | 0,28 | 241 |
| FREM1 | AF-Q5H8C1-F1 | P_0 | 0,65 | 0,81 | 1676,6 | 1888,8 | 31,7 | 0,42 | 252 |
| FREM1 | AF-Q5H8C1-F1 | P_1 | 0,63 | 0,80 | 1223,5 | 1435,1 | 31,3 | 0,37 | 191 |
| FREM1 | AF-Q5H8C1-F1 | P_2 | 0,61 | 0,81 | 1149,4 | 1244,4 | 35,7 | 0,33 | 209 |
| FREM1 | AF-Q5H8C1-F1 | P_4 | 0,63 | 0,83 | 856,4 | 1061,6 | 20,6 | 0,56 | 137 |

**Table S3.** Binding sites identified by DogSiteScorer with a SimpleScore ≥ 0.6 and DrugScore ≥ 0.6. Descriptors: **SimpleScore**, score assigned by the DogSiteScorer used for pocket classification; **DrugScore**, predicted probability that the pocket is druggable; **volume**, geometric volume of the pocket (in Å³); **surface**, surface area of the pocket (in Å²); depth, depth of the pocket (in Å); **HPh**, number of hydrophobic contacts of the pocket; and **siteAtoms**, number of atoms that define the pocket.

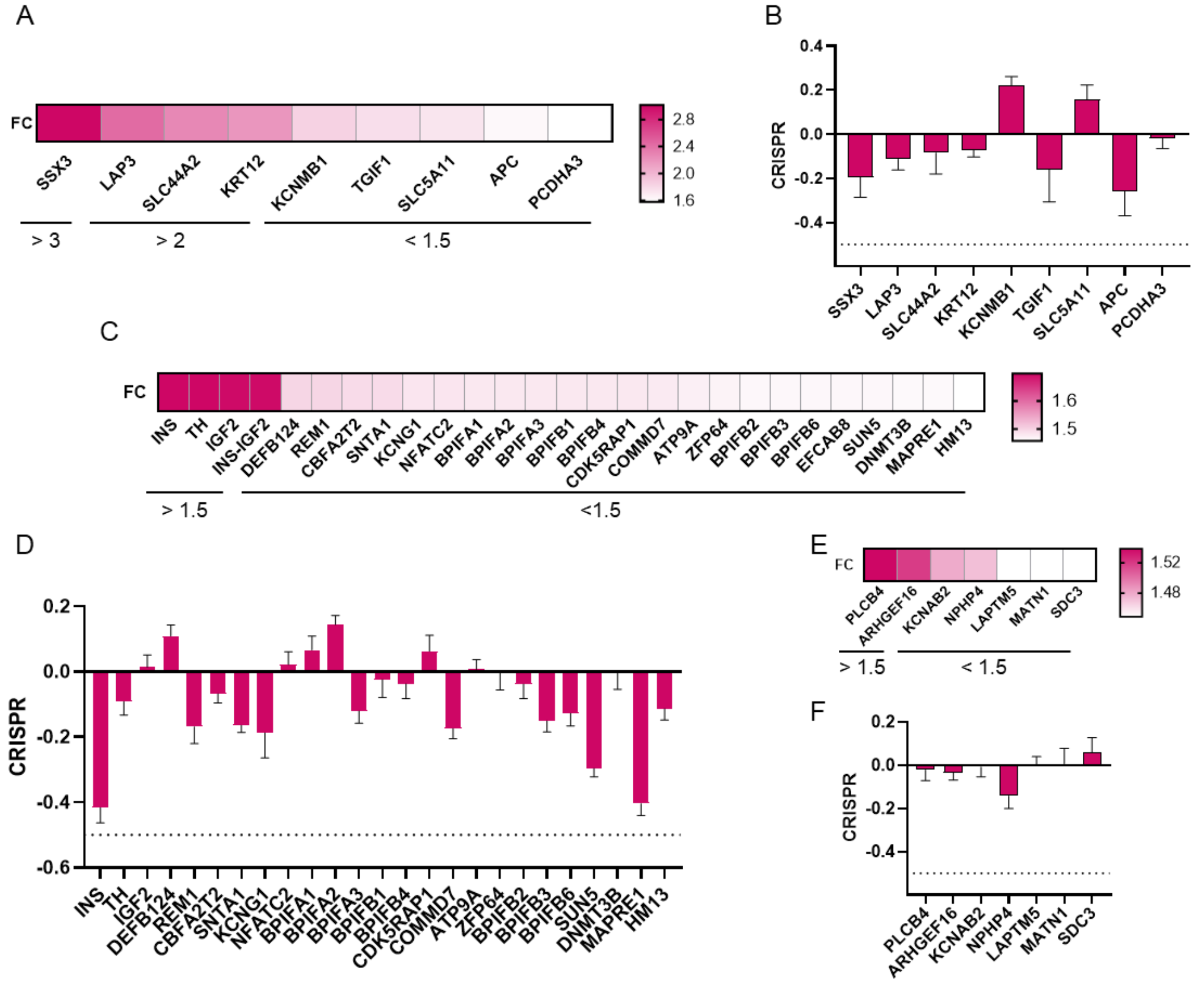


**Figure S1. Fold change and genetic dependency for genes associated with the mutational and copy-number context of LGR5-high colorectal tumours.** Heat maps showing the expression fold change between LGR5-high and LGR5-low COAD tumours for genes associated with (A) somatic mutations, (C) copy-number amplifications and (E) copy-number deletions, ordered by decreasing fold change. (B, D, F) CRISPR gene effect scores (Chronos) for the corresponding genes across colorectal cancer cell lines, as described in Methods; bars show the mean and error bars the [SEM/SD]. The dotted line indicates −0.5, below which a gene is conventionally considered essential. No gene in any set crosses this threshold

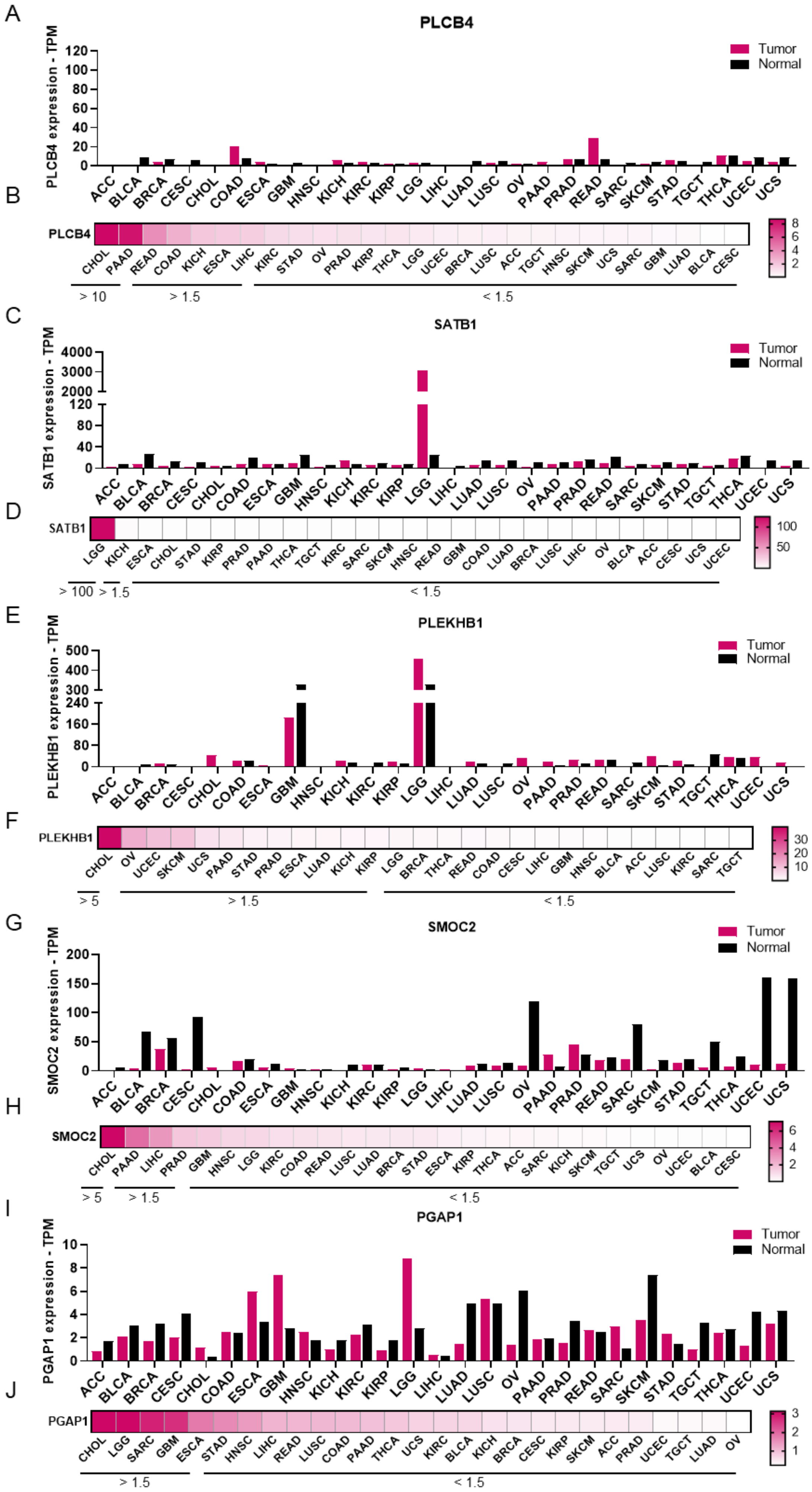
A
PLCB4
Tumor
Normal
PLCB4 expression - TPM
ACC BLCA BRCA CESC CHOL COAD ESCA GBM HNSC KICH KIRC KIRP LGG LIHC LUAD LUSC OV PAAD PRAD READ SARC SKCM STAD TGCT THCA UCEC UCS
B
PLCB4
CHOL PAAD READ COAD KICH ESCA LIHC KIRC STAD OV PRAD KIRP THCA LGG UCEC BRCA LUSC ACC TGCT HNSC SKCM UCS SARC GBM LUAD BLCA CESC
> 10
> 1.5
< 1.5
C
SATB1
Tumor
Normal
SATB1 expression - TPM
D
SATB1
LGG KICH ESCA CHOL STAD KIRP PRAD PAAD THCA TGCT KIRC SARC SKCM HNSC READ GBM COAD LUAD BRCA LUSC LIHC OV BLCA ACC CESC UCS UCEC
> 100
> 1.5
< 1.5
E
PLEKHB1
Tumor
Normal
PLEKHB1 expression - TPM
F
PLEKHB1
CHOL OV UCEC SKCM UCS PAAD STAD PRAD ESCA LUAD KICH KIRP LGG BRCA THCA READ COAD CESC LIHC GBM HNSC BLCA ACC LUSC KIRC SARC TGCT
> 5
> 1.5
< 1.5
G
SMOC2
Tumor
Normal
SMOC2 expression - TPM
H
SMOC2
CHOL PAAD LIHC PRAD GBM HNSC LGG KIRC COAD READ LUSC LUAD BRCA STAD ESCA KIRP THCA ACC SARC KICH SKCM TGCT UCS OV UCEC BLCA CESC
> 5
> 1.5
< 1.5
I
PGAP1
Tumor
Normal
PGAP1 expression - TPM
J
PGAP1
CHOL LGG SARC GBM ESCA STAD HNSC LIHC READ LUSC COAD PAAD THCA UCS KIRC BLCA KICH BRCA CESC KIRP SKCM ACC PRAD UCEC TGCT LUAD OV
> 1.5
< 1.5

**Figure S2. Pan-cancer tumour-versus-normal expression of the five intracellular candidates upregulated in LGR5-high colorectal tumours.** Expression of PLCB4 (A, B), SATB1 (C, D), PLEKHB1 (E, F), SMOC2 (G, H) and PGAP1 (I, J) across TCGA tumour cohorts and normal tissue. (A, C, E, G, I) Median expression in transcripts per million (TPM) in tumour (magenta) and normal (black) tissue for each of the 27 TCGA cohorts, identified by their standard study abbreviations. Y-axis scales are gene-specific and differ between panels to accommodate the wide range of absolute expression across candidates. (B, D, F, H, J) Heat maps of the fold change between tumour and non-transformed tissue for the corresponding gene, with cohorts ordered left to right by fold change; colour scales are gene-specific and are shown to the right of each panel. Horizontal bars beneath each heat map group cohorts by ratio band, as indicated.

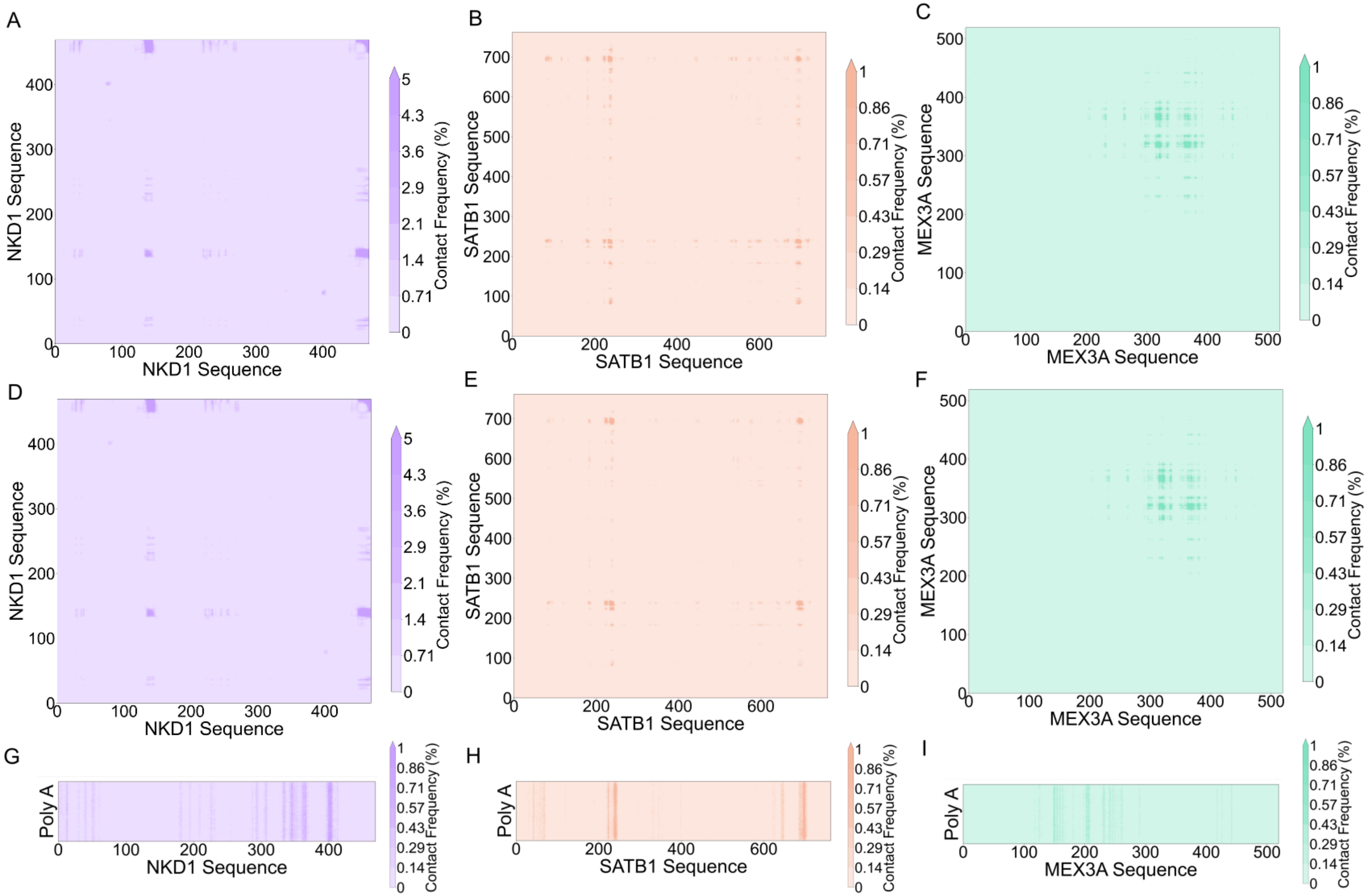


**Figure S3. Residue-resolved intermolecular contact maps for NKD1, SATB1, and MEX3A in the absence and presence of poly(A) RNA. (A-C)** Contact frequency maps for the homotypic protein-protein interface (NKD1-NKD1, SATB1-SATB1, and MEX3A-MEX3A, respectively) in the protein-only systems. **(D-F)** Contact frequency maps for the same homotypic interfaces in the corresponding protein + poly(A) systems. **(G-I)** Contact frequency maps between each protein and poly(A) RNA in the protein + poly(A) systems. Color scales indicate contact frequency (%) and are shown independently for each panel.

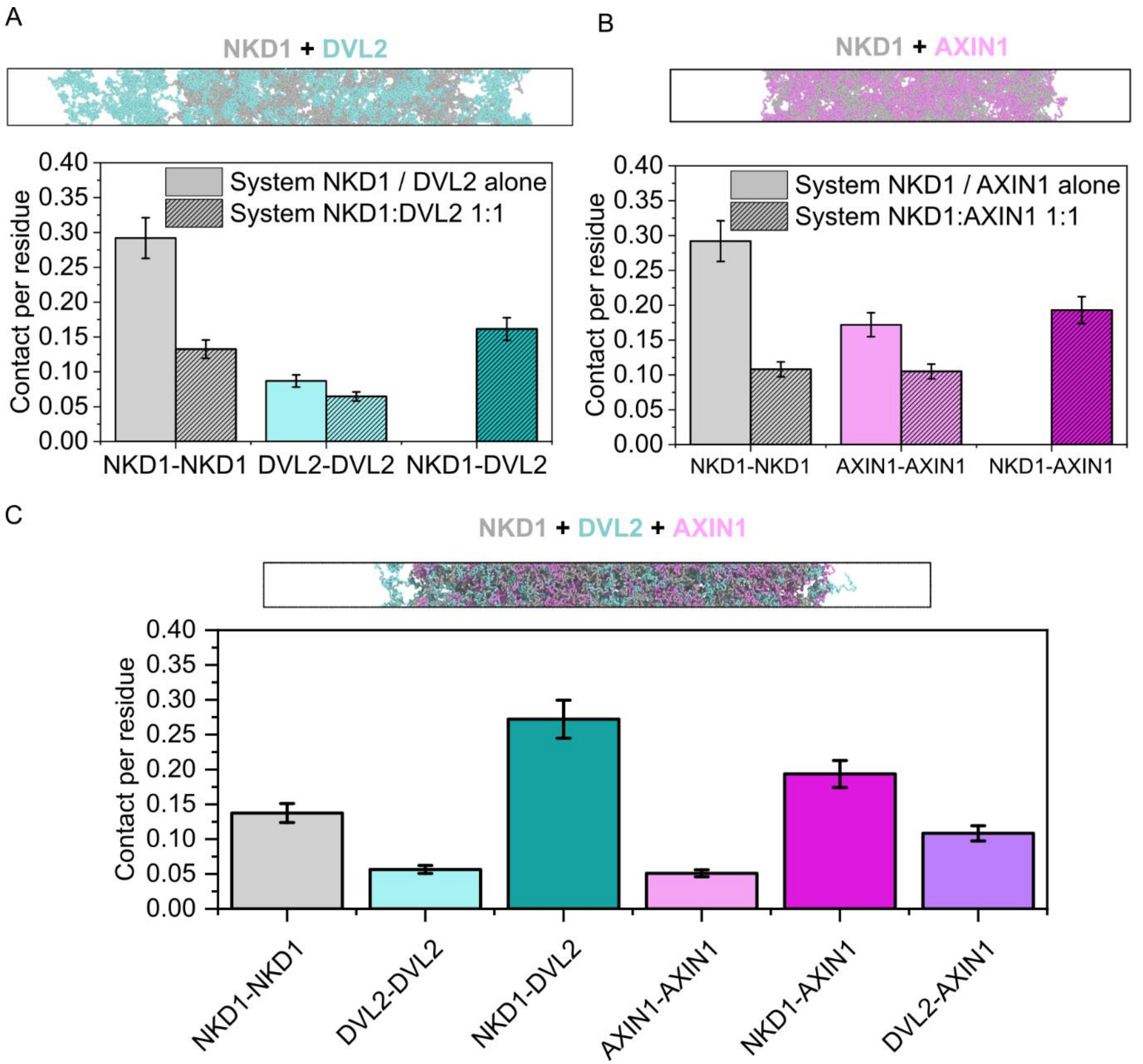


**Figure S4. Intermolecular contacts per residue for the NKD1-DVL2-Wnt system and its binary subsystems. (A)** Contacts per residue for NKD1-NKD1, DVL2-DVL2, and NKD1-DVL2 pairs, comparing NKD1 and DVL2 simulated alone (solid bars) versus the binary NKD1:DVL2 (1:1) system (hatched bars). **(B)** Contacts per residue for NKD1-NKD1, AXIN1- AXIN1, and NKD1- AXIN1 pairs, comparing NKD1 and AXIN1 simulated alone (solid bars) versus the binary NKD1: AXIN1 (1:1) system (hatched bars). **(C)** Contacts per residue for all homotypic and heterotypic pairs (NKD1-NKD1, DVL2-DVL2, NKD1-DVL2, AXIN1- AXIN1, NKD1- AXIN1, and DVL2- AXIN1) in the ternary NKD1:DVL2:AXIN1 (2:1:1) system. Error bars represent standard error of the mean.

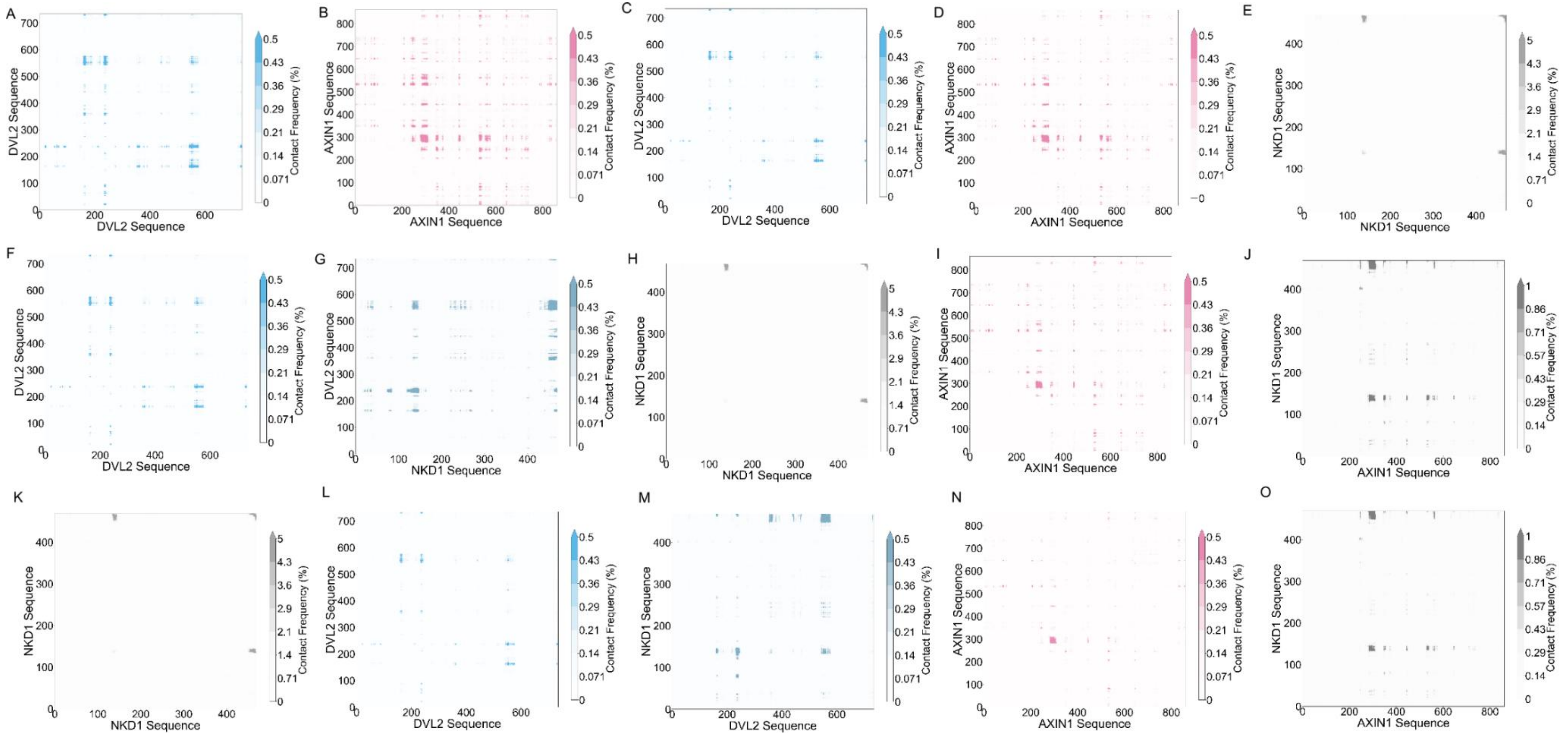


**Figure S5. Residue-resolved intermolecular contact maps across the NKD1, DVL2, and AXIN1 simulated systems. (A)** DVL2-DVL2 contacts in the DVL2-only system. **(B)** AXIN1- AXIN1 contacts in the AXIN1-only system. **(C-D)** DVL2-DVL2 and AXIN1- AXIN1 contacts, respectively, in the binary DVL2: AXIN1 (1:1) system. **(E-G)** NKD1-NKD1, DVL2-DVL2, and NKD1-DVL2 contacts, respectively, in the binary NKD1:DVL2 (1:1) system. **(H-J)** NKD1-NKD1, AXIN1- AXIN1, and NKD1- AXIN1 contacts, respectively, in the binary NKD1: AXIN1 (1:1) system. **(K-O)** DVL2-DVL2, NKD1-NKD1, AXIN1- AXIN1, DVL2- AXIN1, and NKD1- AXIN1 contacts, respectively, in the ternary NKD1:DVL2: AXIN1 (2:1:1) system. Color scales indicate contact frequency (%) and are shown independently for each panel.
